# Parvalbumin basket cell myelination accumulates axonal mitochondria to internodes

**DOI:** 10.1101/2022.04.07.487496

**Authors:** Koen Kole, Bas J. B. Voesenek, Maria E. Brinia, Naomi Petersen, Maarten H. P. Kole

## Abstract

Parvalbumin-expressing (PV^+^) basket cells are fast-spiking interneurons that exert critical control over local neuronal circuit activity and oscillations. PV^+^ interneuron axons are partially myelinated but the electrical and metabolic roles of myelin in axonal functions remain poorly understood. Here, we developed Cre-dependent AAV vectors for cell type-specific investigation of mitochondria with genetically encoded fluorescent probes. Single-cell reconstructions and mining of ultrastructural data revealed that mitochondria selectively cluster to myelinated segments of PV^+^ basket cell axons. Cuprizone-induced demyelination abolished mitochondrial clustering in PV^+^ axons but increased axonal mitochondrial densities in excitatory axons. The internodal clustering of mitochondria was preserved with genetic deletion of myelin basic protein, suggesting that noncompacted myelin is sufficient. Finally, two-photon imaging of action potential-evoked mitochondrial calcium (mt-Ca^2+^) responses showed that internodal mitochondria did not contribute in buffering activity-dependent Ca^2+^ influx. These findings suggest that oligodendrocyte-PV^+^ axon signaling assembles mitochondria to branch selectively fine-tune metabolic demands.

## Introduction

In the vertebrate central nervous system, oligodendrocytes wrap multilamellar membranes around axons to form myelin sheaths, which reduce local internodal depolarization, speed up action potential propagation and enhance action potential fidelity^1,2^. Although the myelin sheath strongly facilitates the electrical conductivity of internodes, the insulation also prevents the axolemma from taking up glucose and other metabolites from the extracellular space. Glycolytic oligodendrocytes aid neurons in overcoming this hurdle by providing trophic support to the axon via the myelin sheath in the form of pyruvate and lactate^3,4^, and by modulating the function of axonal mitochondria via exosomes^5^. Mitochondria are versatile organelles that are chiefly committed to the production of adenosine triphosphate (ATP), and in axons require metabolic support from the myelin sheath in order to preserve their integrity^6^. Previous research, mostly performed in white matter tracts and glutamatergic axons, has shown that demyelination causes dysfunction and shape changes of axonal mitochondria in both multiple sclerosis (MS)^7–9^ and experimental models^10–15^. For example, the mitochondrial size and density increase together with electron transport chain protein expression, and mitochondria lose their membrane potential which is critical for ATP synthesis^7,8,11,14^. The accumulation of mitochondria upon demyelination, together with a low mitochondrial content in myelinated axon and vice versa^16–19^, have given rise to the general notion that myelination reduces the need for mitochondria.

Myelin is also present around inhibitory GABAergic axons, and in the neocortex nearly exclusively those belonging to parvalbumin-expressing (PV^+^) basket cells^20–22^. Comprising the most abundant interneuron cell type in the cortex, PV^+^ basket cells are characterized by a highly complex and extensive axonal organization enabling critical control of the local circuitry and over gamma oscillations^23,24^. These interneurons possess small diameter axons (∼0.4 μm, ref^25^) which could make them highly vulnerable to pathology of the myelin sheath^26,27^. We recently found that experimental demyelination through toxic or genetic means critically disrupts inhibitory transmission by PV^+^ interneurons, characterized by fewer presynaptic terminals, reduced intrinsic excitability and an abolishment of gamma oscillations^22,28^. Interestingly, postmortem studies show that MS is associated with a specific loss in PV^+^ interneurons and their presynaptic terminals^28–30^. With a low threshold of action potential generation, sustained high firing frequencies and synchronization with fast gamma oscillations, PV^+^ interneurons use large amounts of energy to fuel the sodium/potassium ATPase activity to maintain the balance of ionic gradients^31,32^. In line with their high energy usage, PV^+^ interneurons are characterized by a relatively high mitochondrial content and cytochrome c oxidase^33,34^ while deficits in the spatial distribution of mitochondria in PV^+^ axons are associated with impairments of gamma oscillations^35^. One leading hypothesis is that myelination provides trophic support to mitochondria in PV^+^ axons to maintain their levels of high-frequency spike generation and activity during gamma oscillations ^21,36,37^.

To examine how myelination affects mitochondrial distribution and function we developed a Cre-dependent viral approach to express mitochondria-targeted genetically encoded fluorescent reporters, mt-GFP and mt-GCamp6f in Cre-driver lines. Two-photon guided patch-clamp recordings from PV^+^ interneurons of the somatosensory cortex followed by confocal microscopy and reconstructions with submicron precision revealed that in normally myelinated axons mitochondria are clustered to myelinated segments of PV^+^ axons, which was disrupted following experimental demyelination. Analyses of a publicly available electron microscopy (EM) data set^38^ confirmed the higher mitochondrial content in myelinated segments of (PV^+^) basket cells but showed homogeneous distribution along partially myelinated layer 2/3 pyramidal neurons. In addition, two-photon imaging of action potential-evoked mitochondrial Ca^2+^ transients (mt-GCamp6f) showed that mitochondria in myelinated internodes displayed low or no Ca^2+^ buffering activity, which increased following demyelination. These results indicate that in neocortical PV^+^ basket cells axons myelination spatially organizes mitochondria, locally increasing the mitochondrial content within internodes, revealing a novel and direct metabolic support by myelination.

## Results

### An AAV-mediated approach for cell type-specific labeling of mitochondria

We first developed a Cre-dependent adeno-associated viral (AAV) vector (AAV-EF1a-mt-GFP-DIO; **Figure 1A**) and examined the *in vivo* labelling of mitochondria. The use of an AAV vector allows control over transduction rates to enable single-cell analyses, an important advantage over brain-wide cell type-specific expression^39^. Upon injection into the somatosensory cortex (S1) of either PV-Cre (labelling PV^+^ interneurons) or Rbp4-Cre (labelling cortical layer 5 pyramidal neurons) transgenic mice (**Figure 1B**), this viral vector resulted in GFP-labeled mitochondria in molecularly defined cell types (**Figure 1C-F**). Expression of mt-GFP-DIO in PV-Cre; Ai14 mice (expressing tdTomato in PV^+^ interneurons) labelled interneurons across cortical layers 2 to 6 at high specificity (243/248 cells or 97.94% mt-GFP^+^ cells were tdTomato^+^; **Figure 1C**; ref^40^). Furthermore, two-photon-guided targeted whole-cell recordings from mt-GFP-expressing (mt-GFP^+^) cells revealed that all cells fired action potentials at high frequency (123.41 ± 13.13 Hz upon 500 pA current injection; *n =* 8 cells from 5 mice). Action potentials were brief in half-width (295.74 ± 2.52 μs; *n* = 5 cells from 4 mice) and showed a prominent afterhyperpolarization (−15.83 ± 2.25 mV; *n* = 5 cells from 4 mice, **Figure 1D**), consistent with previously described properties of neocortical PV^+^ interneurons^32^.

**Figure 1.**
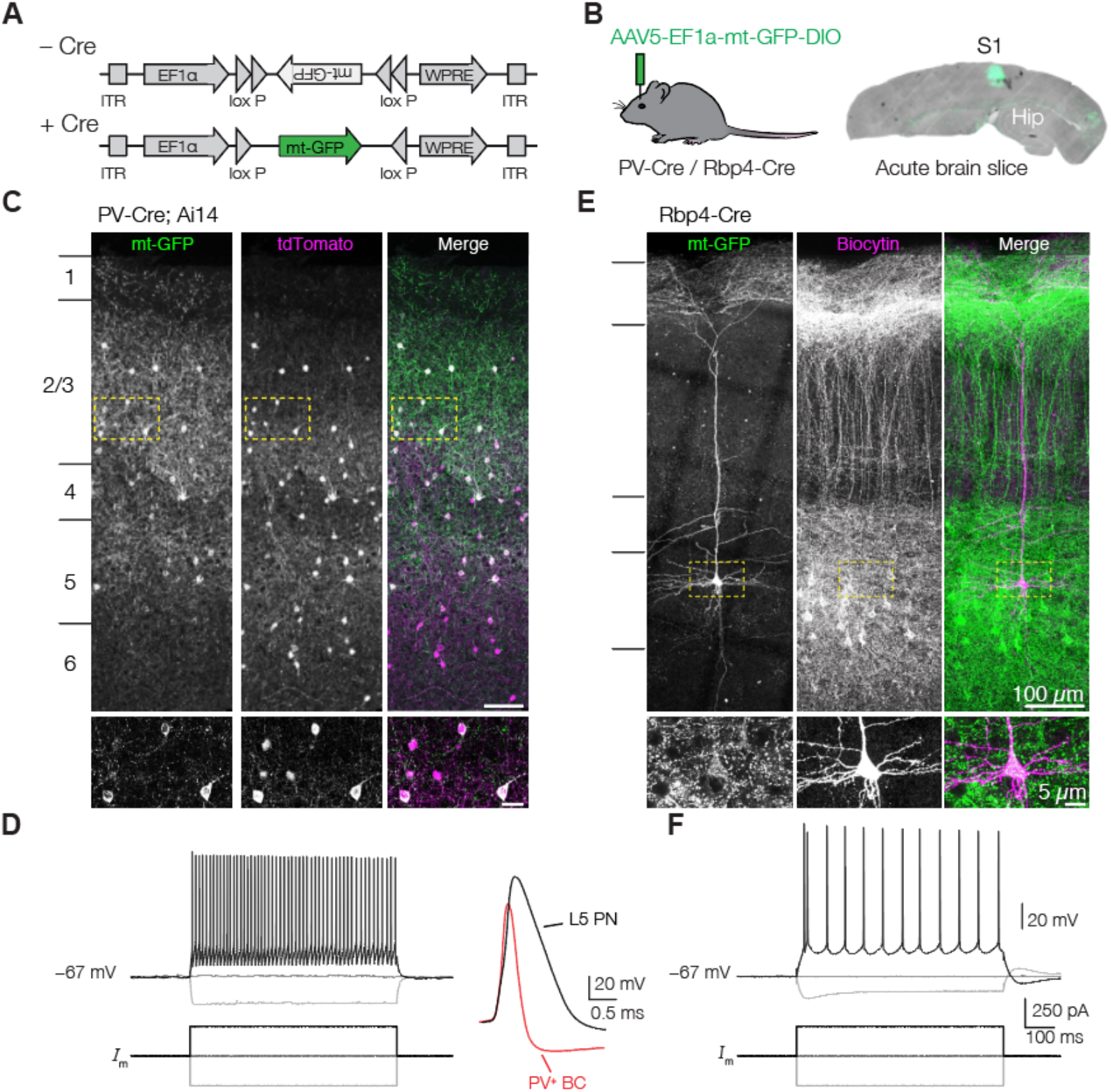
mt-GFP expresses in a Cre-dependent manner. **(A)** Schematic of the AAV construct design. Mitochondria-targeted green fluorescent protein (mt-GFP) requires Cre recombination for expression under the EF1α promoter. **(B)** PV-Cre; Ai14 or Rbp4-Cre mice were injected with AAV5-EF1a-mt-GFP-DIO. Hip, hippocampus. **(C)** Expression in PV-Cre; Ai14 mice results in high overlap between tdTomato and mt-GFP. Dashed boxes indicate area depicted by insets at bottom. **(D)** *Left:* Example traces of action potentials in a mt-GFP^+^ fast spiking PV^+^ basket cell (BC); *Right*: single action potentials in mt-GFP^+^ cells from a PV-Cre; Ai14 or Rbp4-Cre mouse (aligned at onset). **(E)** AAV expression in Rbp4-Cre mice results in mitochondrial labeling in L5 PNs. Note the characteristic apical dendrites visualized by mt-GFP. Dashed boxes indicate area depicted by insets at bottom. **f** Example traces of a current-clamp recording and action potential train in a mt-GFP^+^ L5 PN.

To examine the Cre dependency of the mt-GFP-DIO AAV vector, we expressed mt-GFP^+^ in Rbp4-Cre mice, a Cre-driver line commonly used to label cortical layer (L)5 pyramidal neurons (PNs)^41^. In Rbp4-Cre mice, mt-GFP^+^ somata were pyramidally shaped, restricted to L5 and possessed a large apical dendrite characteristic of L5 PNs (**Figure 1E**; ref.^42^). Mt-GFP^+^ cells in Rbp4-Cre mice always discharged action potentials in low-frequency trains (40.45 ± 4.76 Hz upon 500 pA current injection; *n =* 13 cells from 4 mice) and of comparatively large half width (680.85 ± 15.80 μs; *n =* 11 cells from 4 mice), consistent with the known electrophysiological properties of L5 PN (**Figure 1D, F**; ref.^43^). Mitochondria are motile organelles but are substantially more stationary in adult mice and *in vivo*^44^. We quantified mitochondrial motility in a subset of our mt-GFP^+^ acutely prepared slices and found that the vast majority (∼99%) of mitochondria were stable in both cell types (**Supplementary Figure 1**). Together, these findings indicate that mt-GFP-DIO enables the study of mitochondria in a highly cell-type specific manner.

### Mitochondria are clustered at the axonal initial segment (AIS) and enlarged upon demyelination

To examine the contribution of myelination to mitochondrial distribution we injected PV-Cre; Ai4 mice with AAV-EF1a-mt-GFP-DIO in S1 and fed mice either with control food or a diet supplemented with 0.2% cuprizone, which selectively kills oligodendrocytes, leading to widespread cortical demyelination^45^. We first focused on the axon initial segment (AIS), a highly excitable domain at the base of the axon where action potentials (APs) are initiated and which is known to adapt to demyelination^43,46^. The AIS was identified by immunostaining for the anchoring protein βIV spectrin in cortical tissue from mt-GFP-expressing PV-Cre; Ai14 mice. By contouring of the perimeter of mt-GFP signals we subsequently estimated the shape and size of mitochondria. The results showed that the mitochondrial area was significantly increased in axons from cuprizone-treated mice (nested t-test, *P =* 0.0241; **Figure 2A-C**). In contrast, the mitochondrial aspect ratio (length/width) and density were unaffected (**Figure 2D-E**).

**Figure 2.**
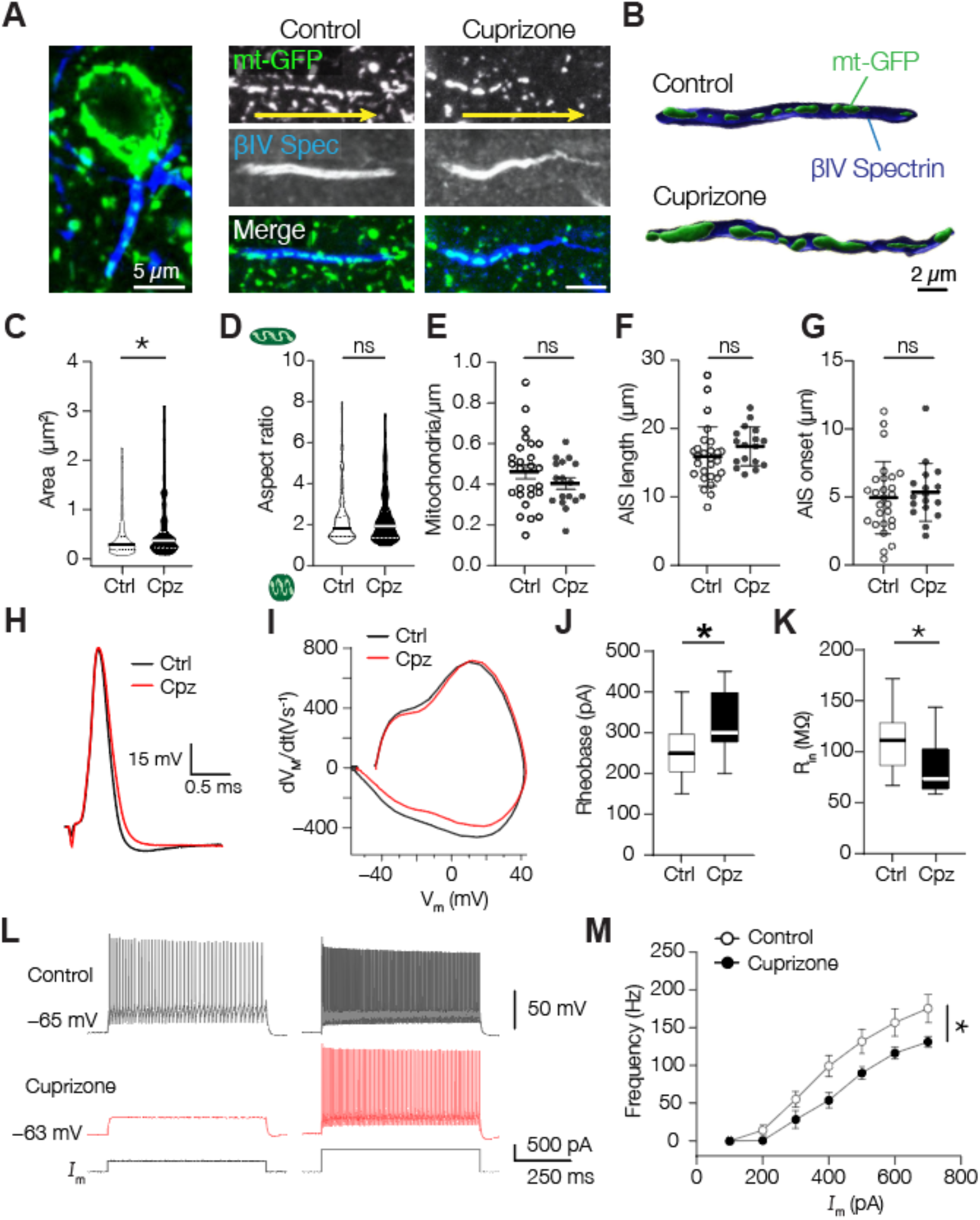
Cuprizone-induced demyelination enlarges AIS mitochondria and reduces PV^+^ interneuron excitability. **(A)** Example images of mitochondria (green) at the AIS (blue) of PV^+^ interneurons. Arrows (yellow) indicate anterograde direction of the axon. **(B)** 3D surface rendering of PV^+^ AISs containing mitochondria of control or cuprizone-treated axons. **(C)** Mitochondrial area is increased upon demyelination (nested t-test, \**P =* 0.0241; df = 43; Ctrl, *n =* 215 mitochondria in 27 AISs from 4 mice; Cpz, *n =* 144 mitochondria in 18 AISs from 3 mice). **(D)** Mitochondrial aspect ratio is unaffected (nested t-test, *P =* 0.5656; df = 43; Ctrl, *n =* 215 mitochondria in 27 AISs from 4 mice; Cpz, *n =* 144 mitochondria in 18 AISs from 3 mice). **(E)** Density of AIS mitochondria is unchanged upon demyelination (unpaired t-test, *P =* 0.2396; df = 41; Ctrl, *n =* 27 AISs from 4 mice; Cpz, *n =* 17 AISs from 3 mice). **(F)** AIS length is unaffected by cuprizone-treatment (unpaired t-test, *P =* 0.2112; df = 0.42; Ctrl, *n =* 27 AISs from 4 mice; Cpz, *n =* 17 AISs from 3 mice). **(G)** AIS distance (soma edge to AIS) is comparable between control and cuprizone-treated groups (unpaired t-test, *P =* 0.6055; df = 42; Ctrl, *n =* 27 AISs from 4 mice; Cpz, *n =* 17 AISs from 3 mice). **(H)** Example single APs of a control and demyelinated PV^+^ interneuron upon a 3 ms current injection. **(I)** Phase-plane plots of the traces shown in d; Reduced PV^+^ interneuron excitability suggested by **(J)** increased rheobase (unpaired t-test, \**P =* 0.0104; df = 33; Ctrl, *n =* 22 cells from 12 mice; Cpz, *n =* 13 cells from 7 mice) and **(K)** reduced input resistance (unpaired t-test, \**P =* 0.0337; df = 33; Ctrl, *n =* 22 cells from 12 mice; Cpz, *n =* 13 cells from 7 mice). **(L)** Example traces of PV^+^ interneuron responses after control (black) or cuprizone (red) treatment in response to 400 (left) or 600 pA (right) current injection. **(M)** Population data showing a reduced firing frequency upon somatic current injection in demyelinated PV^+^ interneurons (two-way ANOVA with Bonferroni’s post hoc test; interaction effect, *P =* 0.0215, F_(6, 126)_ = 2.582; treatment effect \**P =* 0.0439, F_(1, 21)_ = 4.596; Ctrl, *n =* 17 cells from 12 mice; Cpz, *n =* 11 cells from 7 mice). In truncated violin plots (c, d), solid lines represent the median, dotted lines represent 25^th^ and 75^th^ quartiles. Scatter plots (e-g) indicate the mean, individual data points represent cells, error bars indicate SEM; Box plots (j, k) represent the 25^th^ to 75^th^ percentiles, whiskers indicate the maximal and minimal values, solid line represents the median.

Both the AIS length and distance relative to the soma were comparable between groups (**Figure 2F-G**). AIS length and location are critical to the current threshold for AP generation^47^. To test whether AP initiation was preserved we performed whole-cell recordings from PV^+^ interneurons in acute slices. In all slice experiments, we recorded features of excitability in artificial cerebral spinal fluid containing 5 mM L-lactate and 10 mM D-glucose, resembling the *in vivo* extracellular environment containing lactate and low amounts of glucose^48^. In accordance with the unaffected geometrical properties of the AIS, electrophysiological recordings of PV^+^ interneurons revealed no change in AP wave form, voltage or current threshold, amplitude, half-width, or resting membrane potential (**Figure 2H-I**; **Supplementary Figure 2**). However, demyelinated PV^+^ interneurons displayed a higher rheobase (**Figure 2J**), a lower input resistance (**Figure 2K**), and lower firing rate upon sustained somatic current injection (**Figure 2L-M**) indicating reduced excitability, in line with previous recordings^22^. Together, these results show that following demyelination PV^+^ interneuron excitability is reduced, and AIS mitochondria are larger.

### Mitochondria of the demyelinated PV^+^ axon are increased in size and lost in proximal branches

To examine the mitochondrial morphology and distribution throughout the entire cytoarchitecture we reconstructed mt-GFP^+^ axons and dendrites of whole-cell recorded biocytin-filled PV^+^ interneurons (**Supplementary Figs. 3, 4**). Axonal mitochondria in PV^+^ interneurons were often found at branch points and *en passant* boutons and were smaller in appearance compared to their dendritic counterparts (**Supplementary Figure 3**; ref.^49^). Quantification showed that compared to dendritic mitochondria those in axons covered significantly less length, were smaller and more spherical (**Supplementary Figure 3**). These morphological differences between mitochondria are in agreement with previous findings in L2/3 PNs ^50^ and the observed mitochondrial properties in L5 PNs (**Supplementary Figure 4**).

Consistent with our observations at the AIS, demyelination significantly increased the size of axonal mitochondria (**Figure 3A-B**), with a mean surface area increase of ∼60% (nested t-test, *P =* 0.044; **Figure 3B-C**), without affecting mitochondrial shape (**Figure 3D**), suggesting a uniform increase. Importantly, PV^+^ interneuron myelination is most abundant at lower axonal branch orders (≤ 4^th^ order)^20,21,25^. To test whether demyelination affects axonal mitochondria differently we separated the distal population from the 2^nd^ to 4^th^ branch orders and found that the demyelination-induced mitochondrial size increase was comparable (**Figure 3E**).

**Figure 3.**
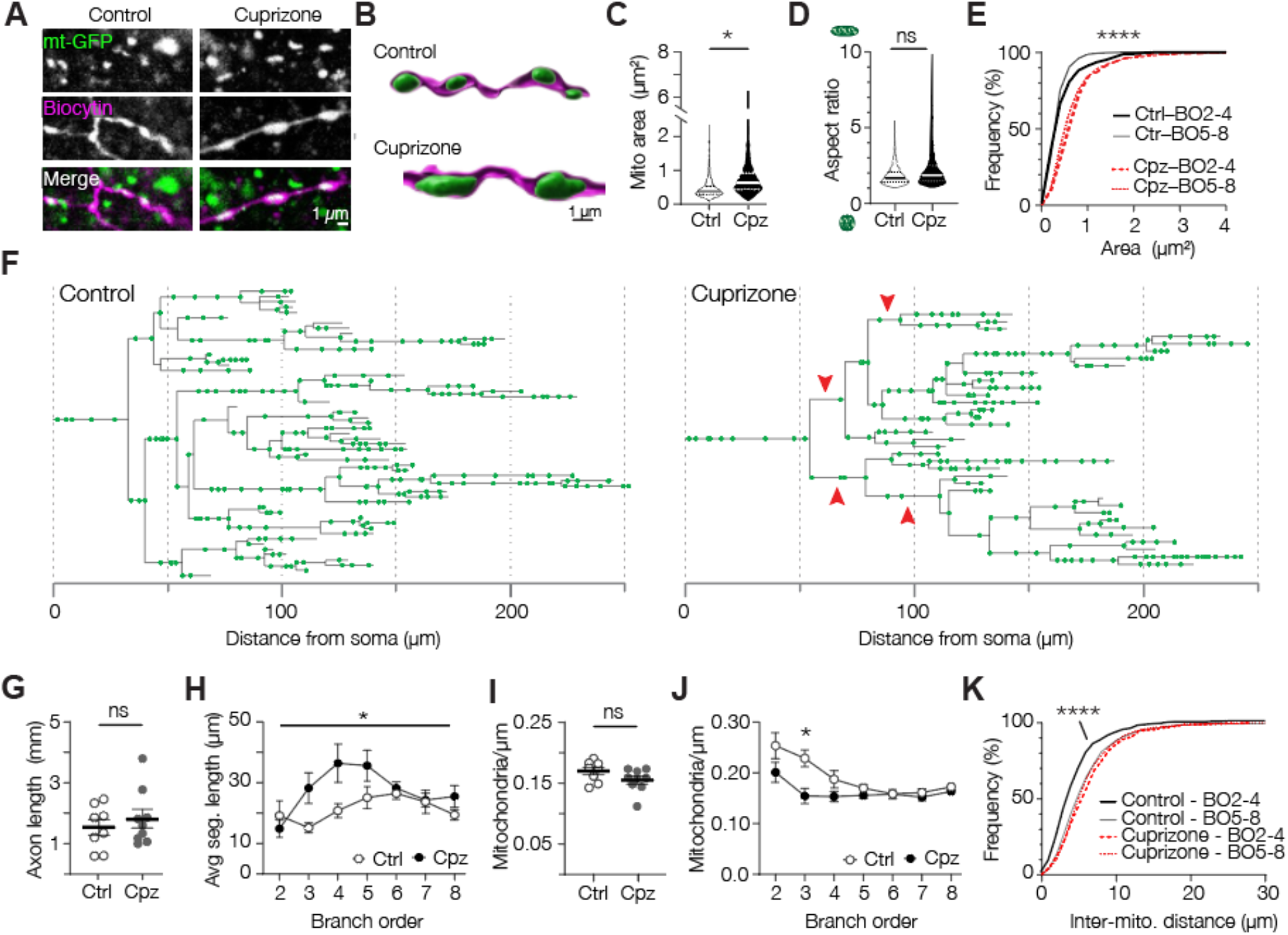
Cuprizone-induced demyelination reduces mitochondrial density in proximal PV^+^ interneuron axons. **(A)** Example confocal images of mitochondria (mt-GFP) in control (left) and demyelinated axon (right). **(B)** 3D rendered images of mt-GFP and biocytin. **(C)** Increased mitochondrial size upon demyelination (nested t-test, \**P =* 0.038; df = 8; Ctrl, *n =* 5 axons, 805 mitochondria; 5 cells from 3 mice, Cpz, *n =* 5 axons, 1218 mitochondria; 5 cells from 4 mice). **(D)** Unchanged mitochondrial aspect ratio in demyelinated axons (nested t-test, *P =* 0.2962; df = 8; Ctrl, *n =* 5 axons, 805 mitochondria; 5 cells from 3 mice; Cpz, *n =* 5 axons, 1218 mitochondria; 5 cells from 4 mice). **(E)** Cumulative distribution plots of mitochondrial size showing uniform increase across all branch orders (BO, Kruskal-Wallis test \*\*\*\**P* < 0.0001; Dunn’s post hoc test, All Ctrl vs Cpz comparisons, *P* < 0.0001, Ctrl BO 2-4 vs BO 5-8, *P =* 0.8402; Cpz BO 2-4 vs BO 5-8, *P =* 0.3932. Ctrl, *n =* 5 axons, 805 mitochondria; 5 cells from 3 mice. Cpz, *n =* 5 axons, 1218 mitochondria; 5 cells from 4 mice). **(F)** Example axonograms of a control-(*left*) and a cuprizone-treated (*right*) PV^+^ interneuron. Red arrowheads indicate examples of second, third or fourth branch orders where mitochondria appear relatively sparse. **(G)** Cuprizone treatment induces no change in axonal length (unpaired t-test, *P =* 0.4918; df = 15; Ctrl, *n =* 8 axons, 5 mice; Cpz, *n =* 9 axons, 5 mice). **(H)** Increased segment length in demyelinated PV^+^ axons as a function of branch order (two-way ANOVA; branch order x treatment effect, \**P =* 0.0440, F_(6,90)_ = 2.266; branch order effect, *P =* 0.0104, F_(3.916, 58.71)_ = 3.660; treatment effect, *P =* 0.0479, F_(1,15)_ = 4.639; Bonferroni’s post hoc test n.s.; Ctrl, *n =* 8 axons, 5 mice; Cpz, *n =* 9 axons, 5 mice). **(I)** No change in overall axonal mitochondrial density (unpaired t-test, *P =* 0.1170; df = 15; Ctrl, *n =* 8 axons, 5 mice; Cpz, *n =* 9 axons, 5 mice). **(J)** Reduced mitochondrial density in proximal (< 5) but not distal branch orders (two-way ANOVA, Branch order x treatment effect, *P =* 0.0169, F_(6,90)_ = 2.747; branch order effect, *P* < 0.0001, F_(2.764, 41.45)_ = 9.560; treatment effect, *P =* 0.0143, F_(1,15)_ = 7.679; Bonferroni’s post hoc test Branch order 3, \**P =* 0.0340; Ctrl, *n =* 8 axons, 5 mice; Cpz, *n =* 9 axons, 5 mice). **(K)** Cumulative distribution plot showing increased distance between mitochondria in second to fourth branch orders of demyelinated axons but no change in later branch orders (Kruskal-Wallis test *P* < 0.0001; Dunn’s post hoc test, Ctrl vs Cpz BO2-4, \*\*\*\**P* < 0.0001; Ctrl BO2-4 vs Ctrl BO5-8, *P* < 0.0001; Cpz BO2-4 vs Cpz BO5-8, *P =* 0.6008; Ctrl, *n =* 8 axons, 5 mice; Cpz, *n =* 9 axons, 5 mice). Solid lines in truncated violin plots (c, d) represent the median, dotted lines represent 25^th^ and 75^th^ quartiles. Horizontal bars (g, i) represent the mean, individual data points represent axons, error bars represent SEM.

The PV axons did not show signs of swellings or degradation and, consistent with previous work^22^, the total reconstructed axonal length was similar between control and demyelinated axons (unpaired t-test, P = 0.4918; **Figure 3G**). Interestingly however, quantification by branch order revealed an increased average segment length in proximal branches of demyelinated PV^+^ axons (**Figure 3H**). The observation of local structural changes, together with previous work showing that demyelination increases mitochondrial density^7,8,12^, prompted us to investigate the mitochondrial density in PV^+^ axons. While the total mitochondrial density across all branches was comparable between control and demyelinated PV^+^ axons (unpaired t-test, *P =* 0.1170; **Figure 3I**), mitochondria in the proximal branch orders of demyelinated axons were on average ∼20% reduced in density (two-way ANOVA, branch order × treatment effect, *P =* 0.0169; **Figure 3J**). In higher branch orders (≥ 5) the mitochondrial densities were similar between groups (two-way ANOVA with Bonferroni’s post hoc test, *P* > 0.9999). In line with these observations, we found an increased inter-mitochondrial distance specifically in proximal branches of demyelinated axons (≤ 4^th^ order; Kruskal-Wallis test with Dunn’s post hoc test, *P* < 0.0001; **Figure 3K**). These results are surprising since previous studies consistently reported mitochondria increase in demyelinated axons^12,13^. To examine mitochondria specifically in glutamatergic L5 axons we used a similar viral strategy in Rbp4-Cre mice and examined mitochondrial density in the white matter below the somatosensory cortex. In contrast to the mitochondria loss in PV^+^ interneurons we found a *higher* density of mitochondria after cuprizone-mediated demyelination (unpaired t-test, *P* = 0.0441; **Supplementary Figure 5**). Together, these results indicate that myelin loss results in a reduced mitochondrial clustering specifically in the proximal and putatively demyelinated branches of the PV^+^ axon.

### Mitochondria are clustered at the myelinated PV^+^ axon

The loss of mitochondrial clustering in proximal axons following demyelination suggests that mitochondrial distribution is determined, at least in part, by the presence of the myelin sheath. If internode myelination is sufficient to cluster mitochondria this should be visible in the proximal axon arbors of normally myelinated PV^+^ interneurons, which are characterized by heterogeneous myelin patterns^20,21,25^. To test this, we performed a triple staining for myelin basic protein (MBP), GFP and biocytin in sections from control mice and reconstructed PV^+^ axons to quantify the mitochondrial density in MBP positive (MBP^+^) and negative (MBP^−^) axonal segments (**Figure 4A-B, Supplementary Figure 6**). In agreement with previous findings^20^, MBP^+^ segments were ∼19 μm in length (on average 18.59 ± 1.84 μm, range: 2.56–47.62 μm, *n =* 32 internodes). Interestingly, axonal mitochondria in myelinated segments were larger (nested t-test, *P =* 0.0121; **Figure 4C**), and more elongated compared to mitochondria in unmyelinated branches (nested t-test, *P =* 0.0388; **Figure 4D**). We observed that MBP^+^ segments contained more mitochondria (paired t-test, *P* = 0.0013, **Figure 4E**), and were also significantly longer than MBP^−^ segments (paired t-test, *P* = 0.0267, **Figure 4F**). Consistent with our hypothesis, after correcting for length, MBP^+^ axonal segments displayed a significantly higher mitochondrial density compared to MBP^−^ segments of the same axon and comparable branch orders (**Figure 4A-B, G**). Furthermore, mitochondrial densities in control MBP^−^ segments were not different from cuprizone-induced demyelinated PV^+^ axons (One-way ANOVA, *P* = 0.0138; Bonferroni’s post hoc test, MBP^−^ vs. MBP^+^ *P* = 0.0356, MBP^+^ vs. cuprizone BO3 *P* = 0.0276; MBP^−^ vs cuprizone BO3 P > 0.9999). These results support the idea that myelin wrapping alone suffices to cluster and increase the size of mitochondria locally within PV^+^ internodes.

**Figure 4.**
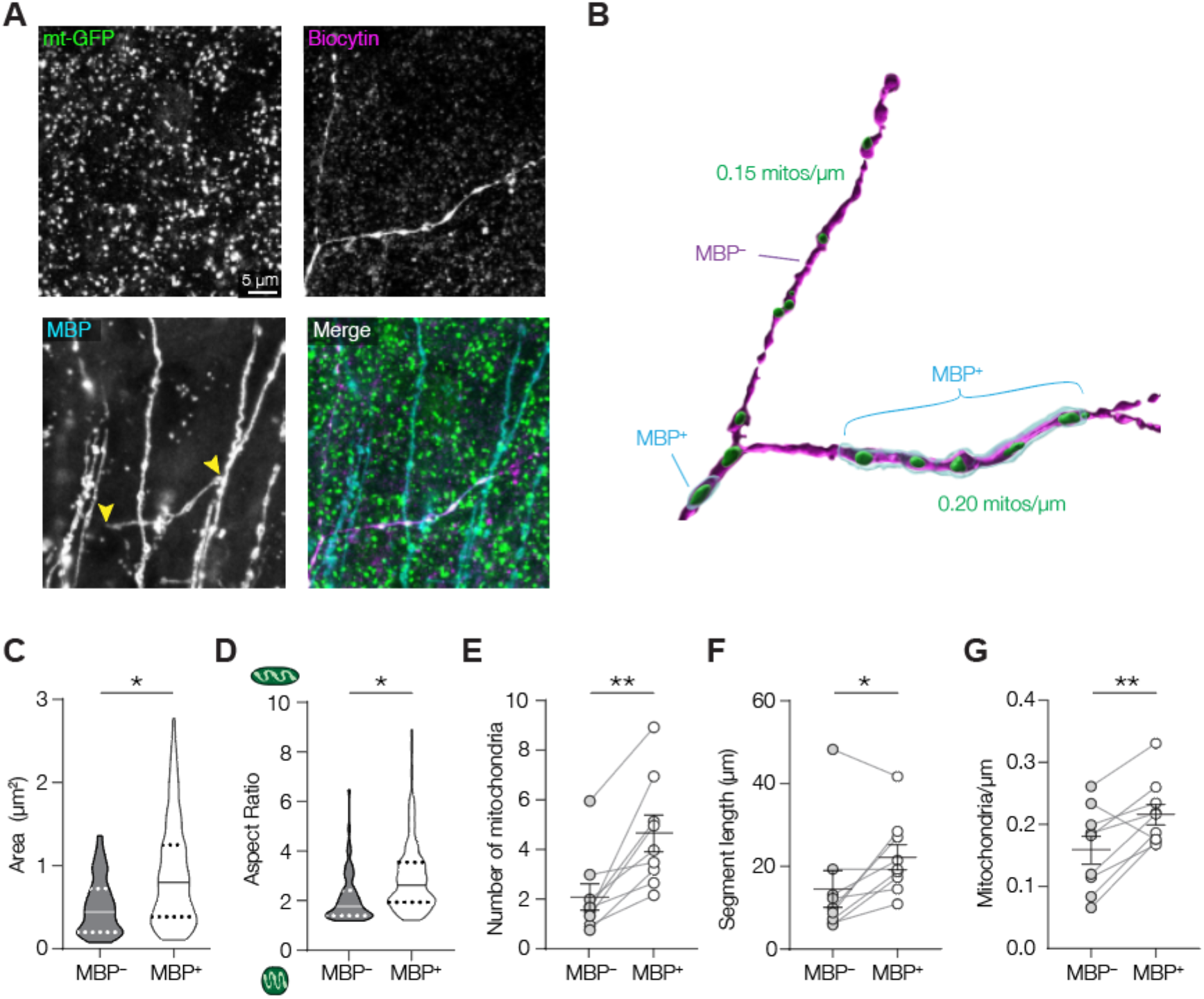
Mitochondria are clustered to myelinated segments of PV^+^ axons. **(A)** Confocal images of a control PV^+^ axon branch point with one myelinated (MBP^+^) and one unmyelinated (MBP^−^) daughter branch. **(B)** 3D surface rendering of the confocal images in a. **(C)** Mitochondrial area is higher at the myelin sheath (nested t-test, *P =* 0.0121; df = 14; unmyelinated, *n =* 68 mitochondria; myelinated, *n =* 114 mitochondria, 8 axons from 6 mice). **(D)** Mitochondrial aspect ratio is significantly higher in myelinated axons (nested t-test, *P =* 0.0388; df = 14; unmyelinated, *n =* 68 mitochondria; myelinated, *n =* 114 mitochondria, 8 axons from 6 mice). **(E)** More mitochondria in MBP^+^ segments (paired t-test,\*\**P* = 0.0013, df = 8; *n =* 9 axons from 6 mice). **(F)** Myelinated PV^+^ axonal segments are significantly longer (paired t-test, \**P* = 0.0267, df = 8; *n =* 9 axons from 6 mice). **(G)** Mitochondrial density in the myelinated segment is higher compared to unmyelinated ones within the same axon (paired t-test, \*\**P* = 0.0099; *n =* 9 axons from 6 mice). Solid lines in truncated violin plots (c, d) represent the median, dotted lines represent 25^th^ and 75^th^ quartiles, individual data points represent axons (1-4 segments per axon).

### Myelination clusters mitochondria selectively to baskets cell internodes

The unexpected axonal mitochondrial clustering within myelinated PV^+^ BC internodes could be a common feature of axon-glia signalling along intermittently myelinated axons. To test this, we used a recently published, fully annotated and minable 3D electron microscopy (EM) dataset of the mouse visual cortex^38^. We identified (putative PV^+^) basket cells based on their morphological properties and presence of myelinated arbours (**Figure 5A**; see Methods) and compared axonal mitochondrial distribution with those of the primary axons of L2/3 PNs which are known to exhibit patchy distribution of myelin along the primary axon^51^ (**Figure 5B**).

**Figure 5.**
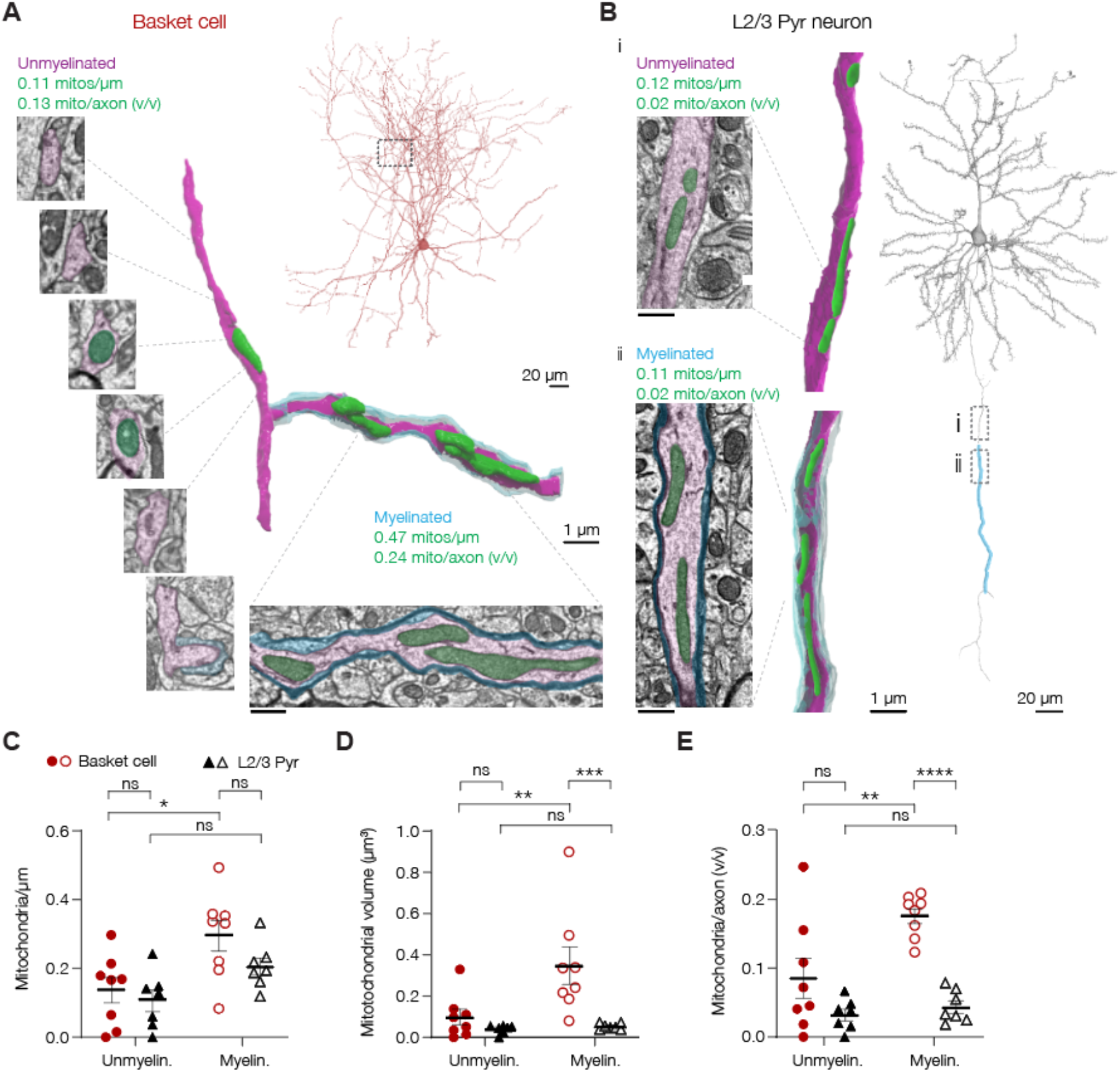
Myelination assembles mitochondria in basket cell-but not PN internodes. **(A-B)** 3D EM renders and reconstructions of a basket cell (A) and L2/3 PN (B) axon with and without myelin. **(C)** Mitochondrial density is increased in myelinated basket but not L2/3 PN segments (two-way ANOVA, myelination x cell type effect, *P =* 0.4162, F_(1,13)_ = 0.7052; myelination effect, *P* = 0.0034, F_(1,13)_ = 12.82; cell type effect, *P =* 0.1206, F_(1,13)_ = 2.760; Bonferroni’s post hoc test, basket myelin. vs. unmyelin., \**P =* 0.0130; PN myelin. vs. unmyelin., *P =* 0.1665; Myelin. basket vs PN, *P =* 0.1766; Unmyelin. basket vs PN *P* > 0.9999). **(D)** Mitochondria in basket myelinated axonal segments are larger compared to those in PNs (two-way ANOVA, myelination x cell type effect, *P =* 0.0430, F_(1,13)_ = 5.028; myelination effect, *P* = 0.0285, F_(1,13)_ = 6.070; cell type effect, *P =* 0.0043, F_(1,13)_ = 11.89; Bonferroni’s post hoc test, basket myelin. vs. unmyelin., \*\**P =* 0.0087; PN myelinated vs. unmyelin., *P >* 0.9999; Myelin. basket vs PN, \*\*\**P =* 0.0009; Unmyelin. basket vs PN *P* = 0.9090; basket, myelin. *n* = 68, unmyelin. *n* = 18 mitochondria; PN, myelin. *n* = 53, unmyelinated *n* = 21 mitochondria). **(E)** In myelinated basket but not PN segments mitochondria take up a larger fraction of the axonal volume (two-way ANOVA, myelination x cell type effect, *P =* 0.0410, F_(1,13)_ = 5.143; myelination effect, *P* = 0.0117, F_(1,13)_ = 8.578; cell type effect, *P =* 0.0002, F_(1,13)_ = 27.17; Bonferroni’s post hoc test, basket myelin. vs. unmyelin., \*\**P =* 0.0044; PN myelin. vs. unmyelin., *P >* 0.9999; Myelin. basket vs PN, \*\*\*\**P <* 0.0001; Unmyelin. basket vs PN *P* = 0.0837). Scale bars for EM images in a and b represent 0.5 μm. Bars in c-e represent means, data points in c and e represent individual axons (basket, *n* = 8 cells; PN, *n* = 7 cells; 1-4 segments per cell for each cell type), data points in d represent average mitochondrial volume per cell (basket, *n* = 8 cells, *n* = 68 mitochondria in myelinated and 18 mitochondria in unmyelinated segments; PN, *n* = 7 cells; *n* = 21 mitochondria in myelinated and 53 in unmyelinated segments). Solid horizontal bars indicate means, error bars indicate SEM, individual datapoints indicate cells.

Consistent with our immunofluorescence analysis, basket cell myelinated segments were significantly longer (two-way ANOVA with Bonferroni’s post hoc test, *P* = 0.0004, **Supplementary Figure 6**). Within basket cell axons mitochondria were found in ∼60% of branch points with at least one myelinated branch (presumed nodes; **Supplementary Figure 6**). Interestingly, mitochondria typically avoided ∼2 μm regions near the edges of the myelin sheath corresponding to the paranodal loops^17,52^, but were with high probability and high densities distributed within the internode (**Supplementary Figure 6**). Myelinated segments of basket cells, both quantitatively and qualitatively in line with the immunoanalysis (Figure 4), had an increased mitochondrial density compared to unmyelinated ones (two-way ANOVA with Bonferroni’s post hoc test, *P* = 0.0130, **Figure 5C**). Interestingly, mitochondrial density in myelinated internodes of L2/3 PNs was not different compared to upstream or downstream segments that were unmyelinated (two-way ANOVA with Bonferroni’s post hoc test, *P* = 0.1665, **Figure 5C**).

Furthermore, in basket cells and in keeping with our immunofluorescence data (c.f. Figure 4), mitochondria within myelinated segments were significantly larger (*P* = 0.0087, **Figure 5D, Supplementary Figure 6**). Mitochondrial size was, however, not affected in L2/3 PN axonal segments (two-way ANOVA with Bonferroni’s post hoc test, *P* > 0.9999, **Figure5D**). Strikingly, when comparing the mitochondria between the two cell types, those in basket cells were ∼7-fold larger (two-way ANOVA with Bonferroni’s post hoc test, *P* = 0.0009, **Figure 5D**). Consistent with previous reports^25^ we noticed that unmyelinated segments were typically thinner. A lower cytoplasmic volume could reduce the need for mitochondria and explain the lower mitochondrial density and smaller mitochondrial size. However, plotting the ratio of mitochondrial to axonal volume showed that mitochondria in myelinated basket cell axons occupied a significantly larger volume compared to unmyelinated branches (two-way ANOVA with Bonferroni’s post hoc test, *P* = 0.0044, **Figure 5E**). In L2/3 PNs the relative volume of mitochondria was comparable between myelinated and unmyelinated segments (two-way ANOVA with Bonferroni’s post hoc test, *P* > 0.9999, **Figure 5E**).

Taken together, the 3D EM results provide independent support of our AAV immunofluorescence results, revealing with ultrastructural detail that mitochondria selectively cluster to myelinated PV^+^ basket cell internodes. Furthermore, they suggest that the clustering of large mitochondria at myelinated axonal segments is not merely a consequence of patchy myelination but may be specific to PV^+^ basket cells.

### Compact myelin is not required for mitochondrial clustering to myelinated PV^+^ axons

Our data so far suggest that oligodendroglia myelination directly induces mitochondrial clustering at myelinated internodes in PV^+^ axons. Axonal mitochondria are supplied with metabolites from glycolytic oligodendrocytes via a system of cytoplasmic channels within myelin and transported across the adaxonal membrane and axolemma^3,4,53^. To test the role of compact and noncompacted myelin we used the *Shiverer* mouse line. These mice harbour a deletion in the *Mbp* gene severely reducing MBP protein levels and resulting in myelin wrapping with only a few noncompacted oligodendroglial membranes^54,55^. We crossed PV-Cre; Ai14 mice with *Shiverer* mice (PV-Cre; Ai14 × Shiverer) and injected wildtype (*Mbp*^WT^) and homozygous (*Mbp*^*Shi*^) mice with AAV-EF1a-mt-GFP-DIO to target GFP to mitochondria of normal or dysmyelinated PV^+^ interneurons. Patch-clamp recordings from PV^+^ interneurons revealed no differences between *Mbp*^WT^ or *Mbp*^Shi^ mice for any of the electrophysiological properties (**Supplementary Figure 7**). During whole-cell recordings, cells were filled with biocytin after which we performed a triple staining for biocytin, GFP and myelin oligodendrocyte glycoprotein (MOG) to identify the location of noncompacted myelin lamellae (**Figure 6A-C**; **Supplementary Figure 7**).

**Figure 6.**
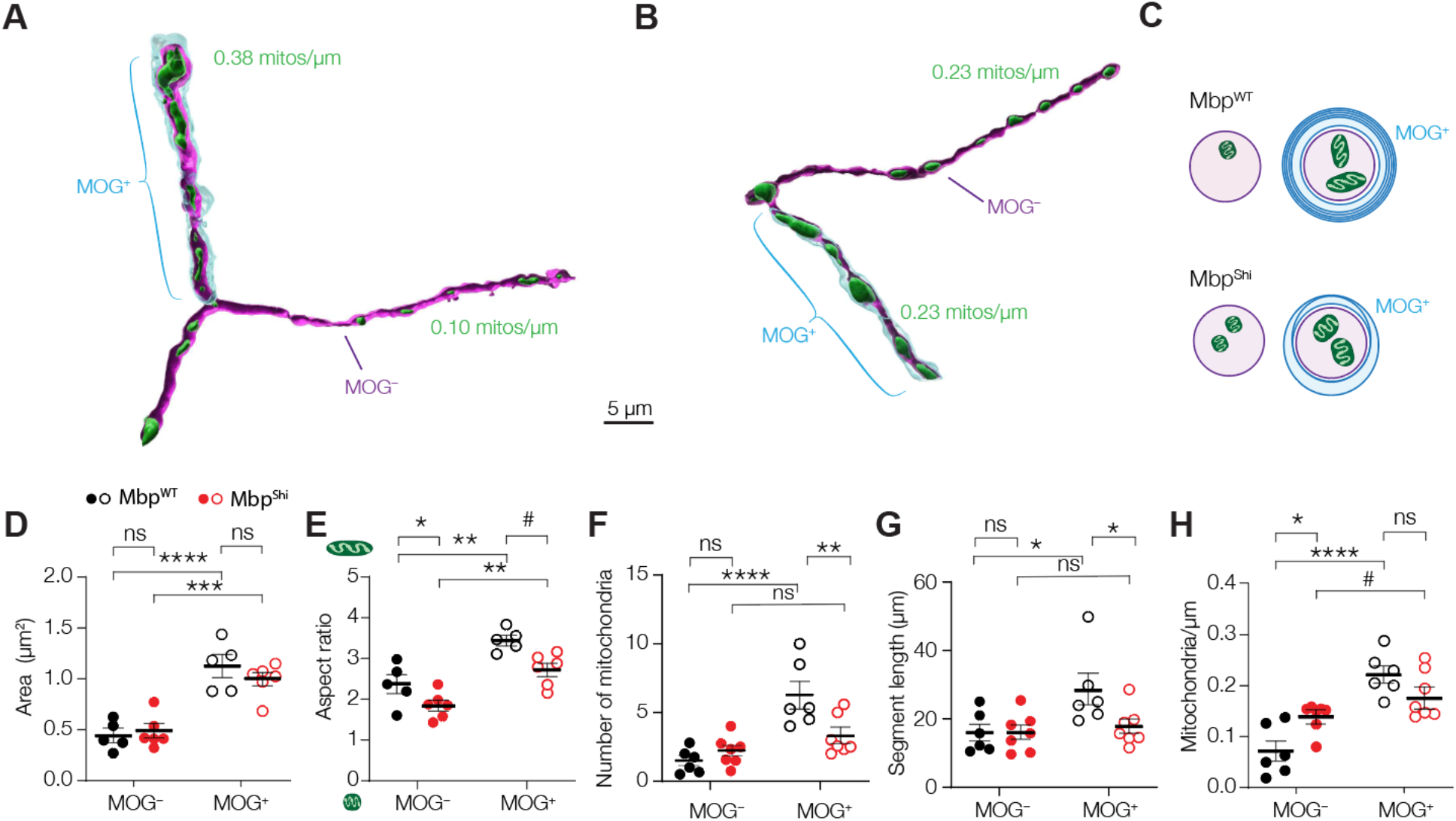
Noncompacted myelin suffices for internodal mitochondrial clustering. **(A-B)** Example 3D renders of a PV^+^ axon of a (A) *Mbp*^*WT*^ or (C) *Mbp*^*Shi*^ mouse (see Supplementary Figure 7). **(C)** Schematic representation of mitochondrial distributions in *Mbp*^*WT*^ and *Mbp*^*Shi*^ PV^+^ axons. **(D)** Mitochondria under myelin sheaths are larger in both *Mbp*^*WT*^ and *Mbp*^*Shi*^ mice (two-way ANOVA, myelination x genotype effect, *P =* 0.1481, F_(1, 9)_ = 2.504; myelination effect, *P <* 0.0001, F_(1, 9)_ = 125.5; genotype effect, *P =* 0.6304, F_(1, 9)_ = 0.2480; Bonferroni’s post hoc test, *Mbp*^*WT*^ MOG^−^ vs. MOG^+^, \*\*\*\**P <* 0.0001; *Mbp*^*Shi*^ MOG^−^ vs. MOG^+^, \*\*\**P <* 0.0001; MOG^−^ *Mbp*^*WT*^ vs. *Mbp*^*Shi*^, *P >* 0.9999; MOG^+^ *Mbp*^*WT*^ vs. *Mbp*^*Shi*^, *P =* 0.4792; *Mbp*^*WT*^, *n* = 6 cells from 3 mice, MOG^+^, *n* = 72 mitochondria, MOG^−^, *n* = 28 mitochondria; *Mbp*^*Shi*^, *n* = 5 cells from 3 mice, MOG^+^, *n* = 78 mitochondria, MOG^−^, *n* = 59 mitochondria). **(E)** Internode mitochondria in both *Mbp*^*WT*^ and *Mbp*^*Shi*^ mice are longer compared to those in unmyelinated axonal segments (two-way ANOVA, myelination x genotype effect, *P =* 0.5428, F_(1, 9)_ = 0.4000; myelination effect, *P =* 0.0002, F_(1, 9)_ = 35.16; genotype effect, *P =* 0.0045, F_(1, 9)_ = 14.13; Bonferroni’s post hoc test, *Mbp*^*WT*^ MOG^−^ vs. MOG^+^, \*\**P =* 0.0032; *Mbp*^*Shi*^ MOG^−^ vs. MOG^+^, \*\**P =* 0.0069; MOG^−^ *Mbp*^*WT*^ vs. *Mbp*^*Shi*^, #*P =* 0.0825; MOG^+^ *Mbp*^*WT*^ vs. *Mbp*^*Shi*^, \**P =* 0.0125; *Mbp*^*WT*^, *n* = 6 cells from 3 mice, MOG^+^, *n* = 72 mitochondria, MOG^−^, *n* = 28 mitochondria; *Mbp*^*Shi*^, *n* = 5 cells from 3 mice, MOG^+^, *n* = 78 mitochondria, MOG^−^, *n* = 59 mitochondria). **(F)** Number of mitochondria in PV^+^ axons of *Mbp*^*WT*^ and *Mbp*^*Shi*^ mice (two-way ANOVA, myelination x genotype effect, *P =* 0.0027, F_(1, 11)_ = 14.77; myelination effect, *P <* 0.0001, F_(1, 11)_ = 37.00; genotype effect, *P =* 0.1406, F_(1, 11)_ = 2.522; Bonferroni’s post hoc test, *Mbp*^*WT*^ MOG^−^ vs. MOG^+^, \*\*\*\**P <* 0.0001; *Mbp*^*Shi*^ MOG^−^ vs. MOG^+^, *P =* 0.2549; MOG^−^ *Mbp*^*WT*^ vs. *Mbp*^*Shi*^, *P =* 0.7802; MOG^+^ *Mbp*^*WT*^ vs. *Mbp*^*Shi*^, \*\**P =* 0.0041; *Mbp*^*WT*^, *n* = 6 cells from 3 mice; *Mbp*^*Shi*^, *n* = 5 cells from 3 mice). **(G)** Segment lengths of PV^+^ interneuron axons in *Mbp*^*WT*^ or *Mbp*^*Shi*^ mice (two-way ANOVA, myelination x genotype effect, *P =* 0.0486, F_(1, 11)_ = 4.914; myelination effect, *P =* 0.0148, F_(1, 11)_ = 8.330; genotype effect, *P =* 0.1348, F_(1, 11)_ = 2.605; Bonferroni’s post hoc test, *Mbp*^*WT*^ MOG^−^ vs. MOG^+^, \**P =* 0.0104; *Mbp*^*Shi*^ MOG^−^ vs. MOG^+^, *P >* 0.9999; MOG^−^ *Mbp*^*WT*^ vs. *Mbp*^*Shi*^, *P >* 0.9999; MOG^+^ *Mbp*^*WT*^ vs. *Mbp*^*Shi*^, \**P =* 0.0315; *Mbp*^*WT*^, *n* = 6 cells from 3 mice; *Mbp*^*Shi*^, *n* = 5 cells from 3 mice). **(H)** Mitochondrial density is higher in internodes in both *Mbp*^*WT*^ and *Mbp*^*Shi*^ mice (two-way ANOVA, myelination x genotype effect, *P =* 0.0034, F_(1, 11)_ = 13.87; myelination effect, *P <* 0.0001, F_(1, 11)_ = 44.39; genotype effect, *P =* 0.4762, F_(1, 11)_ = 0.5441; Bonferroni’s post hoc test, *Mbp*^*WT*^ MOG^−^ vs. MOG^+^, \*\*\*\**P <* 0.0001; *Mbp*^*Shi*^ MOG^−^ vs. MOG^+^, *P =* 0.1070; MOG^−^ *Mbp*^*WT*^ vs. *Mbp*^*Shi*^, \**P =* 0.0168; MOG^+^ *Mbp*^*WT*^ vs. *Mbp*^*Shi*^, *P =* 0.1919; *Mbp*^*WT*^, *n* = 6 cells from 3 mice; *Mbp*^*Shi*^, *n* = 5 cells from 3 mice). Solid horizontal bars indicate means, error bars indicate SEM, individual datapoints indicate cells.

Mitochondria distribution in PV^+^ interneurons from *Mbp*^WT^ mice showed a comparable pattern as found in control mice (**Figure 5, Figure 6**). In brief, we found that mitochondria were both larger and longer in MOG^+^ segments compared to those in MOG^−^ segments (**Figure 6D-E**). We also found more mitochondria in myelinated segments (**Figure 6F**), and myelinated axons were significantly longer than their unmyelinated counterparts (**Figure 6G**). Plotting mitochondria per unit length showed that myelinated axons displayed higher mitochondrial densities compared to those lacking myelin (**Figure 6H**).

In the *Mbp*^Shi^ group, mitochondrial morphology was also myelin-dependent with larger and more tubular mitochondria in MOG^+^ segments compared to those in MOG^−^ branches (two-way ANOVA with Bonferroni’s post hoc test; area, *P* = 0.0001, aspect ratio, *P* = 0.0069; **Figure 6D-E**). In contrast, the number of mitochondria in MOG^−^ and MOG^+^ axonal arbours were comparable (two-way ANOVA with Bonferroni’s post hoc test, *P* = 0.2549; **Figure 6F**). Myelinated segments of PV^+^ axons were also shorter in *Mbp*^Shi^ mice (two-way ANOVA with Bonferroni’s post hoc test, *P* = 0.0315; **Figure 6G**), and there was no difference in length between MOG^+^ and MOG^−^ segments (two-way ANOVA with Bonferroni’s post hoc test, *P* > 0.9999; **Figure 6G**). Interestingly, when correcting for segment length, there was no difference in mitochondrial density between myelinated and unmyelinated axons (two-way ANOVA with Bonferroni’s post hoc test, *P* = 0.1070, **Figure 6H**), due to a significantly increased mitochondrial density in the unmyelinated segments (two-way ANOVA with Bonferroni’s post hoc test, *P* = 0.0168, **Figure 6H**). These results provide further evidence that mitochondria are heterogeneously distributed along the proximal arborization of PV^+^ axons as a function of myelination. In addition, the presence of noncompacted membrane layers suffices for mitochondrial assembly at internodes.

### Myelination attenuates mitochondrial Ca^2+^ buffering

The clustering of mitochondria to internodes may be important for the buffering of Ca^2+^ during neuronal activity^50^. Our recent work showed that myelin strongly attenuates but not completely abolishes depolarization of the axolemma enabling activation of internode voltage-gated channels in particular near the node of Ranvier^1^. Indeed, action potential generation has been shown to cause internodal Ca^2+^ influx spreading far from the node of Ranvier^56,57^ but to which extent this holds true for PV^+^ axons and how mitochondria within and outside internodes respond to activity-dependent Ca^2+^ influx is unknown.

To examine mitochondrial buffering of Ca^2+^ in we employed AAV-mediated and Cre-dependent expression of the mitochondrion-targeted, genetically encoded calcium indicator mt-GCaMP6f in PV^+^ interneurons (**Figure 7**). In acute slices of PV-Cre; Ai14 mice we targeted mt-GCaMP6f^+^ PV^+^ interneurons for whole-cell patch-clamp recordings and filled them with Atto594, visualizing the arborization of axons during 2P recordings and enabling targeted mitochondrial imaging (**Figure 7A**). In the experiments we imaged mt-Ca^2+^ fluorescence responses following a train of ∼100 APs (∼143 Hz). In line with previous findings, we observed that during AP trains mitochondria in putative *en passant* boutons of the distal PV^+^ axon showed strong Ca^2+^ responses, which were unaffected upon demyelination (**Supplementary Figure 8**; ref^50,58^). Similarly, the large amplitude of mt-Ca^2+^ transients at the AIS of PV^+^ interneurons, was unchanged after demyelination (**Supplementary Figure 8**). These findings suggest that activity-dependent mt-Ca^2+^ buffering in presynaptic terminals and the AIS of PV^+^ interneurons is preserved after demyelination.

**Figure 7.**
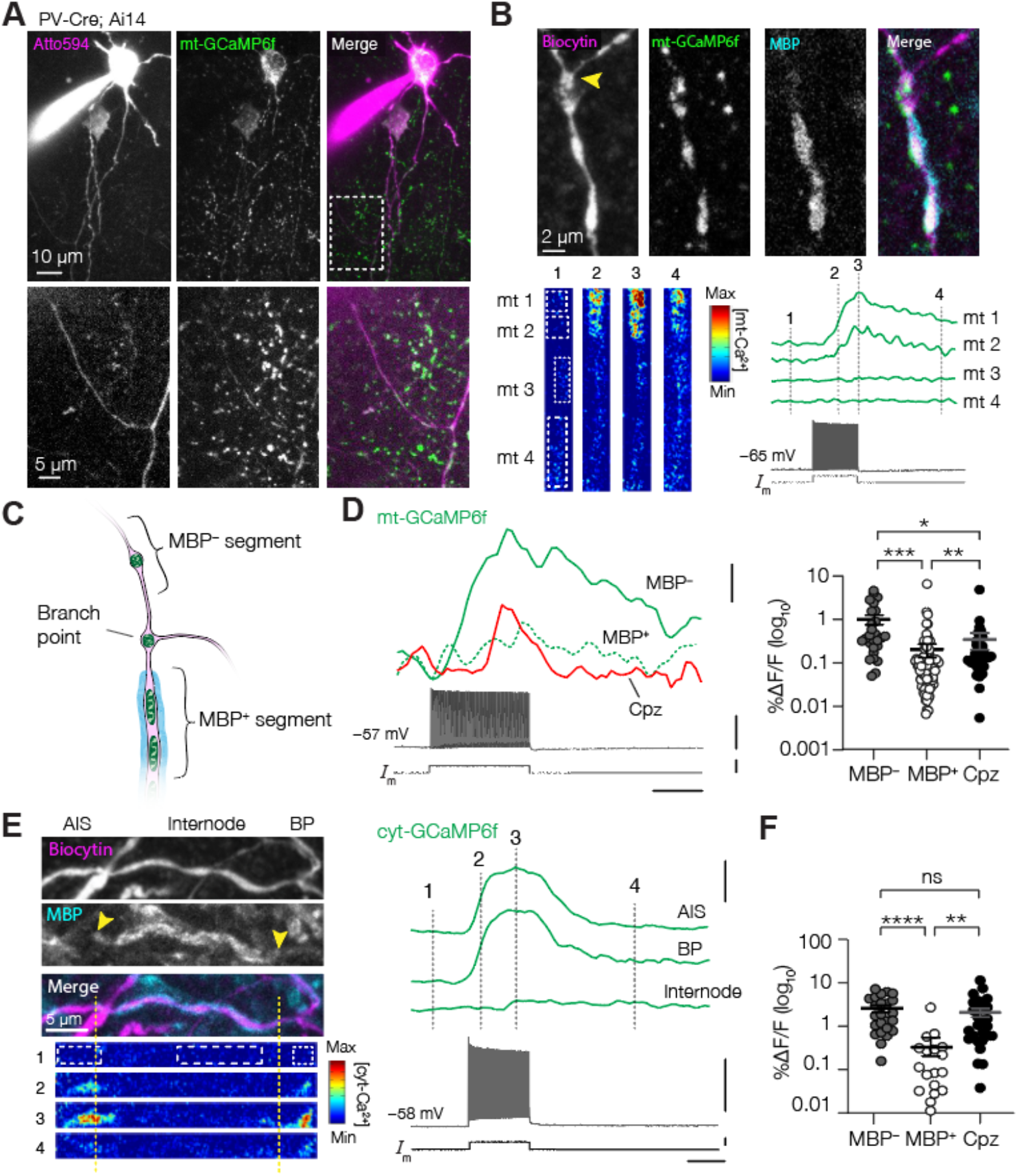
Myelin attenuates internode mt-Ca^2+^ responses. **(A)** Example 2P image of an Atto594-filled mt-GcaMP6f^+^ PV^+^ interneuron. Dashed box indicates the first branch point of the axon, with a characteristic large angle (≥90°). **(B)** Example control PV^+^ axon with mt-GCaMP6f-labeled mitochondria. Mt-Ca^2+^ responses are stronger in MBP^−^ mitochondria (mito 1 and 2) compared to those in MBP^+^ segment (Mito 3 and 4). Dashed lines indicate timepoints corresponding to the heatmap, yellow arrowhead indicates branch point. Scale bars indicate (top to bottom) 100% ΔF/F, 50 mV, 0.5 nA and 0.5 s. **(C)** Schematic overview of sampled axonal regions. In d, for cuprizone, axonal segments in between branch points were selected. **(D)** *Left*, example Ca^2+^ responses of mitochondria in MBP^+^, MBP^−^ or putatively demyelinated segments. *Right*, stronger mitochondrial Ca^2+^ responses in MBP^−^ segments compared to MBP^+^ axon and increased Ca^2+^ responses after demyelination (Kruskal-Wallis test, *P* < 0.0001; Dunn’s post hoc test MBP^−^ vs., MBP^+^ \*\*\**P* < 0.0001; MBP^−^ vs. cuprizone, \**P* = 0.0195; MBP^+^ vs. cuprizone, \*\**P* = 0.0039; MBP^+^, *n =* 101 mitochondria of 12 cells from 7 mice; MBP^−^, *n =* 27 mitochondria of 10 cells from 6 mice; cuprizone, *n* = 36 mitochondria of 10 cells from 5 mice). Scale bars indicate (top to bottom) 25% ΔF/F, 50 mV, 1 nA and 0.5 s. **(E)** *Top:* Example confocal image of the AIS, myelinated internode and first branch point of a PV^+^ axon *Bottom*: time lapse of a two-photon cytosolic (cyt-)GCaMP6f recording of the same axon in response to ∼100 APs. Notice only a slight Ca^2+^ response at the MBP edge but practically none in the rest of the internode. Scale bars indicate (top to bottom) 100% ΔF/F, 50 mV, 0.5 nA and 0.5 s. **(F)** Cyt-Ca^2+^ responses are stronger in unmyelinated and demyelinated axonal segments (Kruskal-Wallis test, P < 0.0001; Dunn’s post hoc test MBP^−^ vs MBP^+^, *P* < 0.0001, MBP^+^ vs cuprizone, *P* = 0.0004, MBP^−^ vs cuprizone, *P* = 0.4351; MBP^+^, *n* = 18 segments of 6 cells from 3 mice; MBP^−^, *n* = 24 segments of 8 cells from 3 mice; cuprizone, *n* = 28 segments of 6 cells from 3 mice). Solid bars in d and f represent the mean, individual data points in d represent mitochondria, individual data points in f represent segments.

To test whether there are differences in internodes, unmyelinated and/or demyelinated branches we imaged mt-GCaMP6f along the proximal arbors (≤5^th^ branch order) and post-hoc immunostained the slices for GFP, MBP and biocytin, allowing unequivocal assessment of the location of mitochondria with respect to myelin sheaths (**Figure 7B-D**). We first focused on segments (*i*.*e*. excluding branch points) and found that mitochondria in control axons showed AP-evoked Ca^2+^ responses with greater amplitudes within MBP^−^ segments compared to those in internodes (Kruskal-Wallis test with Dunn’s post hoc test, *P* < 0.0001; **Figure 7D, Supplementary Video 1**). We next asked whether myelin loss would lead to changes in mt-Ca^2+^ buffering in demyelinated PV^+^ interneurons. When we compared all mt-Ca^2+^ responses in control axons (including both MBP^+^ and MBP^−^ segments) with those in putatively demyelinated axons we found no difference between the two groups (nested t-test, *P =* 0.6055). However, mt-Ca^2+^ transients in cuprizone-treated axons were on average larger compared to MBP^+^ control segments (Kruskal-Wallis test with Dunn’s post hoc test, *P* = 0.0039; **Figure 7D**) but smaller in amplitude compared to MBP^−^ segments (Kruskal-Wallis test with Dunn’s post hoc test, *P* = 0.0195; **Figure 7D**). Mitochondrial Ca^2+^ uptake is known to depend on the influx of extracellular Ca^2+^ (Refs. ^58,59^), which in turn is controlled by myelination^56^. To gain a mechanistic insight into how myelination shapes mt-Ca^2+^, we next examined cytosolic Ca^2+^ (cyt-Ca^2+^) transients in PV^+^ interneurons. To this end, we expressed GCaMP6f in the cytosol of PV^+^ interneurons and imaged the axonal AP-evoked Ca^2+^ transients in MBP^+^, MBP^−^ or putatively demyelinated axonal segments. Similar to our results in mitochondria, cyt-Ca^2+^ transient amplitudes were weak in MBP^+^ arbours (Kruskal-Wallis test with Dunn’s post hoc test, *P* < 0.0001, **Figure 7F, Supplementary Video 2**), while demyelinated axons showed increased Ca^2+^ responses compared to MBP^+^ segments (Kruskal-Wallis test with Dunn’s post hoc test, *P* = 0.0004, **Figure 7F**). These results suggest that myelination strongly attenuates internodal Ca^2+^ entry, limiting downstream mt-Ca^2+^ responses in control PV^+^ axons.

### Myelin loss impairs mitochondrial Ca^2+^ responses at branch points

Upon closer inspection of the localization of the mitochondria with large mt-Ca^2+^ responses we found that they were in close proximity to the outer edge of the myelin sheath or at branch points (**Figure 7B, 8A**). To further identify the precise localization of the myelin borders we immunostained for Caspr, a paranodal marker, which typically had a length of 2.82 ± 0.18 μm (**Figure 8B**; *n =* 36 Caspr^+^ segments of 3 cells from 2 mice). The Caspr^+^ segments often flanked branch points if their branches were myelinated (*n =* 19 branch points of 4 cells from 3 mice, **Figure 8B**). In line with our initial observation, mitochondria were located near Caspr^+^ signals (range 0.01–16.57 μm, on average 2.46 ± 0.96 μm distance, *n =* 20 paranodes of 4 cells from 3 mice; **Figure 8C**). These data suggest that in PV^+^ axons branch points are sites of nodes of Ranvier and mitochondria are often found at the nodal domains (**Supplementary Figure 7**).

**Figure 8.**
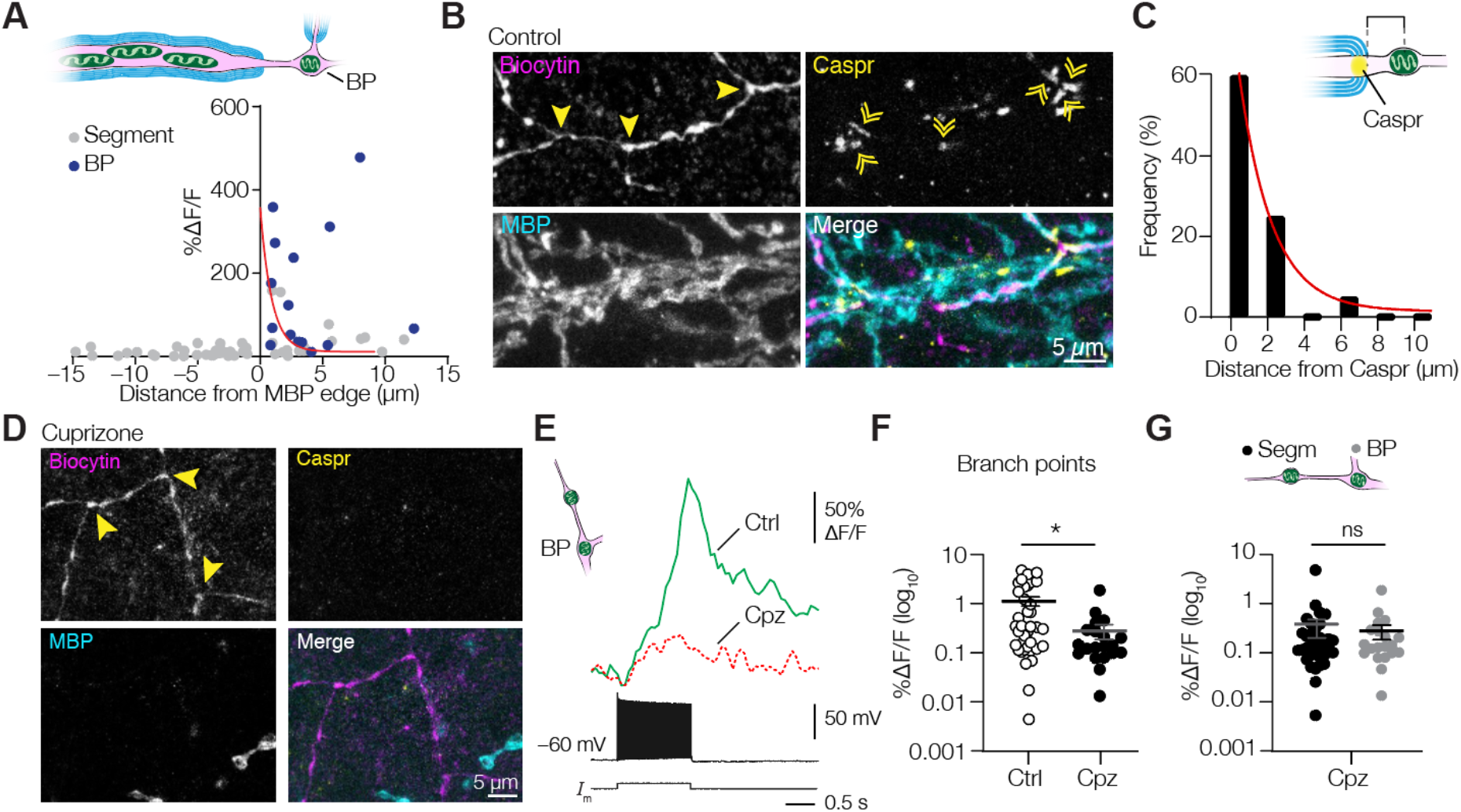
The mt-Ca^2+^ responses are amplified at nodal domains in PV^+^ axons. **(A)** Relation between distances from myelin sheaths and mt-Ca^2+^ response amplitude. Red line indicates exponential decay fit (y = (3.588 – 0.1117) × exp(−0.894 × x) + 0.1117); *n =* 62 mitochondria of 9 cells from 6 mice). **(B)** Example confocal images of PV^+^ axon branch points flanked by paranodes (Caspr). **(C)** Mitochondria are frequently found close to paranodes. Red line indicates exponential decay fit (y = (60.73 – 1.415) × exp(−0.5692 × x) + 1.415); *n =* 20 mitochondria of 4 cells from 3 mice. **(D)** Example of PV^+^ axon branch points of a cuprizone-treated mouse. Notice the overall lack of both MBP and Caspr immunostaining. Solid bars in d and e represent the mean, individual data points represent mitochondria. **(E)** *Left:* schematic representation of the compartments investigated in e and f; *Right*: example traces from control or cuprizone-treated axon branch point mitochondria. **(F)** Amplitude of branch point mt-Ca^2+^ responses was reduced upon demyelination (Mann-Whitney test, \**P* = 0.0123; control, *n =* 38 mitochondria of 11 cells from 7 mice; cuprizone, *n =* 20 mitochondria of 9 cells from 5 mice). **(G)** In demyelinated PV^+^ axons, mt-Ca^2+^ responses at branch points are indistinguishable from those in segments (Mann-Whitney test, *P* = 0.8106; segment, *n* = 36 mitochondria of 10 cells from 5 mice, same data as Figure 7d; branch points, *n =* 20 mitochondria of 9 cells from 5 mice, same data as in f). BP, branch point. Horizontal bars indicate means, error bars indicate SEM.

Finally, demyelination is associated with a disassembly of proteins of the node of Ranvier^43,60,61^. In PV^+^ axons cuprizone-induced demyelination caused a complete loss of Caspr^+^ segments flanking branch points (0 Caspr^+^ segments in 4 cells from 3 animals; **Figure 8D**). To investigate how the nodal mt-Ca^2+^ buffering was affected we compared branch point mt-Ca^2+^ transients^42,65^. The results showed that the AP-evoked mt-Ca^2+^ transients in branch points of demyelinated PV^+^ axons were lower in amplitude (Mann-Whitney test, *P* = 0.0123; **Figure 8E-F**). Moreover, in putatively demyelinated PV^+^ axons, mt-Ca^2+^ responses in branch points were indistinguishable from those in unmyelinated axonal segments (Mann-Whitney test, *P* = 0.8106, **Figure 8G**). Similar to the mt-Ca^2+^ responses, we observed that cyt-Ca^2+^ transients at branch points were reduced upon demyelination (Mann-Whitney test, *P* = 0.0307, **Supplementary Figure 8**). Together, these results suggest that cytoplasmic Ca^2+^ influx determines mt-Ca^2+^ buffering, and that the spatiotemporal distribution with hot spots clustered to branch points and low mt-Ca^2+^ responses within the internodes, is disrupted by demyelination (**Figure 9**).

**Figure 9.**
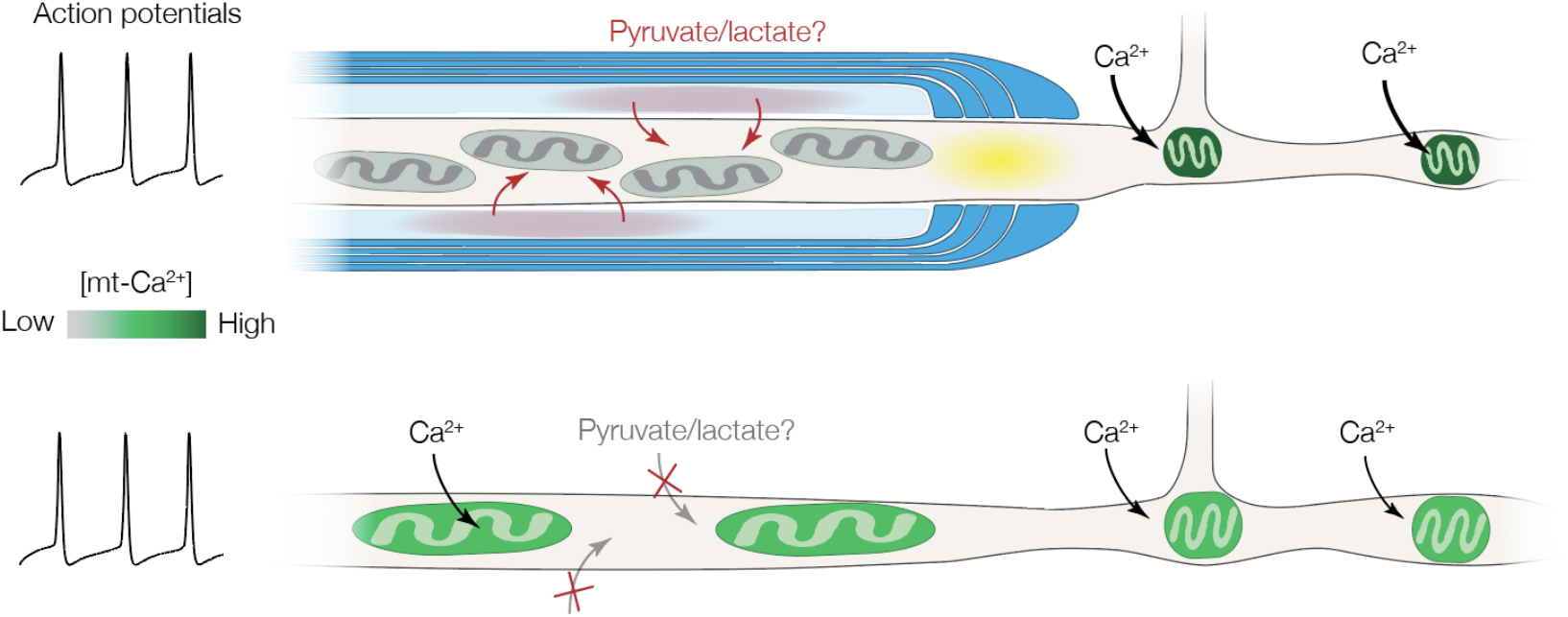
Summary of the effects of myelin loss on mitochondria in PV^+^ axons. In normally myelinated PV^+^ axons (top), mitochondria are clustered at high densities to myelinated internodes (blue), possibly due to local delivery of nutrients such as pyruvate or lactate (red) via cytoplasmic spaces (light blue). Internodal mitochondria avoid the first ∼2 μm of the myelin sheath (paranodal domain, yellow), are relatively large, and display no or very weak activity-dependent mt-Ca^2+^ transients (grey). In branch points and unmyelinated segments, mitochondria are smaller but AP-evoked mt-Ca^2+^ transients are large (dark green). Following myelin loss (bottom), mitochondria become larger and sparser at previously myelinated segments, resulting in a more uniform distribution. The mt-Ca^2+^ responses are uniform and low in amplitude (light green) and nodes of Ranvier are lost.

## Discussion

We used a Cre recombinase mediated viral approach combined with single-cell reconstructions and found that myelinated segments of PV^+^ axons have a high mitochondrial content (**Figure 4, 5**). The locally increased number of internodal mitochondria is in striking contrast with the general notion that myelinated axons contain fewer mitochondria^16–19^. In a previous study based on saturated ultrastructural reconstructions of deep layers of the mouse neocortex it was estimated that the mitochondrial content in myelinated axons was a ∼30-fold lower compared to unmyelinated axons^19^. However, in that study there was no cell type-specific differentiation. Using a publicly available database^38^ we found that the mitochondrial distribution in intermittently myelinated axons differed between (PV^+^) basket cell interneurons and L2/3 PNs. In contrast to the mitochondrial heterogeneity in interneuron axons, axonal mitochondria are homogeneously distributed across myelinated and unmyelinated segments in excitatory axons. In addition, while demyelination reduced mitochondrial density in PV^+^ axons it increased mitochondria in excitatory axons in the white matter (**Fig S5**). The latter observation is consistent with a wealth of data indicating that mitochondrial content increases in experimental demyelination and multiple sclerosis^7,10–13^.

Which factors underlie the cell-type specific clustering of axonal mitochondria in PV^+^ axons remains to be further examined. One possible explanation is that PV^+^ basket cell axons are small in diameter; on average ∼0.3 μm for unmyelinated and ∼0.6 μm for myelinated segments^25,62^. Small diameter axons possess a relatively large membrane surface/cytoplasm ratio limiting axial conductivity for AP propagation but may also require higher energetic costs for axonal transport^63,64^ and impede metabolic supply. The energetic demand of the small diameter PV^+^ interneuron axons is further augmented by the high-frequency firing rates and their generation of gamma frequencies^33^. Taken together, it is tempting to speculate that PV^+^ basket cell axon mitochondria efficiently consume the energy supplied by oligodendrocytes. The myelin sheath and axons of GABAergic neurons show multiple distinctive features in their molecular and cellular organization, including the expression of mitochondrial proteins (reviewed in ^65^). The hypothesis of a direct external metabolic supply is supported by higher levels of 2’,3’-cyclic nucleotide 2’-phosphohydrolase (CNPase)^37^. CNPase is part of the cytoplasmic channels in myelin and the noncompacted inner cytoplasmic loops, involved in lactate and pyruvate supply to the periaxonal space, where these nutrients are shuttled into the axon via monocarboxylate transporters (MCTs)^3,4,6,37^. Indeed, our findings in *Shiverer* mice indicate that noncompacted myelin sufficed to cluster mitochondria at myelin sheaths. Whether oligodendroglial lactate acts as a trophic factor to immobilize axonal mitochondria remains to be tested. Interestingly, high glucose availability has been shown to negatively regulate mitochondrial motility^66^. Furthermore, live imaging in zebrafish axons showed that near paranodes there is myelin-dependent vesicle release^67,68^, a process which requires high levels of ATP^69^. ATP consumption by vesicle release, the Na^+^/K^+^ ATPase^70^ or axonal transport^71^ might cause high local cytoplasmic adenosine diphosphate (ADP) levels, which are known to reduce mitochondrial motility^72^ and thereby may potentially cluster mitochondria to PV^+^ internodes. Future experiments could image ATP within the internodal domains to test the contribution of internodal mitochondria to local ATP synthesis.

To determine the features of motility, Ca^2+^ buffering, anatomical properties of size and shape as well as the distribution of the mitochondria in PV^+^ interneurons we used two-photon and confocal microscopy together with 3D EM data, which have been used previously to study mitochondria^7,9,35,50,73^. Confocal imaging has the advantage of allowing the mapping of mitochondria in large parts of single cells from multiple animals but with the tradeoff of a lower spatial resolution. Therefore, we used 3D EM data and could independently verify at the ultrastructural level that mitochondria are larger and more densely distributed at the myelinated PV^+^ axon, showing the reliability of our confocal approach. Moreover, the ultrastructural data enabled us to directly investigate mitochondria in different cell types from the same brain. We cannot exclude the possibility that cuprizone affects PV^+^ interneuron mitochondria directly, but the increase in mitochondrial size upon demyelination (**Figure 3**) is in keeping with previous studies in axons of other cell types and models of demyelination^7,8,11^.

Our cyt-Ca^2+^ data shows that the diverse mt-Ca^2+^ responses in PV^+^ axons reflect local differentiation of the axon membrane. In normally myelinated axons we observed strong mt-Ca^2+^ influx near branch points which were flanked by Caspr^+^ signals, suggesting that branch points are sites of nodes of Ranvier, consistent with previous work in GABAergic axons^65,74^. At the nodes APs are regenerated causing large local cytoplasmic Ca^2+^ responses as recorded in other neuron types^56,57,75^. The lower Ca^2+^ transients at these sites in both the cytosol and mitochondria after demyelination might therefore reflect loss of nodal ion channel proteins and/or frequency-dependent failures during our trains of ∼100 spikes. These scenarios could be examined in the future using patch-clamp recordings from demyelinated PV^+^ axons to achieve a temporal resolution of single APs. Our cyt-Ca^2+^ imaging data indicates that the large mt-Ca^2+^ transients at unmyelinated segments in control axons are caused by large depolarizations that are accompanied by a strong Ca^2+^ influx. It would be informative to use voltage sensitive dye imaging to examine the contribution of axolemmal depolarization^76,77^. Mitochondria in unmyelinated segments may be located near presynaptic GABAergic terminals where voltage-gated Ca^2+^ channels drive major cytoplasmic Ca^2+^ influx^50,58^. However, mt-Ca^2+^ transients near boutons typically show very large amplitudes (**Supplementary Figure 4**) which we did not observe in MBP^−^ segments.

The present finding that myelin clusters mitochondria and defines mitochondrial Ca^2+^ buffering (which also drives ATP synthesis^78^) indicates that myelination patterns may provide an axoglial mechanism to locally regulate energy homeostasis in the PV^+^ axon arborization. An important caveat is that with our acute slice preparation the myelination state of the axonal segments prior to demyelination is unknown. Future studies could address this by performing longitudinal *in vivo* imaging. Our AAV approach will be useful to deliver vectors that enable monitoring cytosolic ATP in a cell type-specific manner^79^ which will allow the study of axonal ATP levels in (de)myelinated PV^+^ interneurons. Our results also have implications for interneuron myelin plasticity^80,81^. Both myelination and proper mitochondrial distribution in PV^+^ interneurons are known to be important for gamma oscillations, indicating converging pathways that underly PV^+^ interneuron function^22,35^. If in PV^+^ interneurons newly formed myelin sheaths attract mitochondria this could represent a novel axoglial mechanism to selectively alter specific axonal paths in the PV^+^ axonal trees, subsequently strengthening fast inhibitory transmission of specific neocortical connections. Future research will be necessary to test this hypothesis, by for instance using longitudinal mitochondrial imaging during development of myelination or activity-induced myelination. Finally, the present findings may shed light on the vulnerability of PV^+^ basket cells to myelin loss in MS^28–30^. If mitochondria in PV^+^ interneuron axons are particularly reliant on the myelin sheath for external metabolic support, myelin loss might acutely and negatively influence axonal transport, ion homeostasis and other critical processes in PV^+^ axons. Moreover, the small diameter of basket cell axons could make them more vulnerable to excess Ca^2+^ influx upon demyelination^82^, in particular given that mt-Ca^2+^ buffering is insufficient to maintain normal cyt-Ca^2+^ levels during AP trains (**Figure 7**). Together, the data presented here reveal a critical role for myelin to diversify mitochondrial distribution and function in PV^+^ axons and encourage future research into mitochondrial dysfunction in PV^+^ interneuron axons in MS.

## Methods

### Animals

All procedures were performed after evaluation by the Royal Netherlands Academy of Arts and Sciences (KNAW) Animal Ethics Committee (DEC) and Central Authority for Scientific Procedures on Animals (CCD, license AVD8010020172426). The specific experimental designs were evaluated and monitored by the Animal Welfare Body (IvD, protocols NIN19.21.01, NIN19.21.09 and NIN19.21.12). Male and female PV-Cre; Ai14, Rbp4-Cre or Rbp4-Cre; ChETA mice were used in this study that were kept at a 12h day-night cycle with access to food *ad libitum*. At 8 weeks of age, mice were fed either food with or without 0.2% cuprizone (biscyclohexane oxaldihydrazone, Sigma-Aldrich) supplement (range: 5-6 weeks). During this time, the weight and overall welfare of the mice was closely monitored.

### Plasmids

Cre-dependent mt-GFP and mt-GCaMP6f AAV plasmids was created by replacing the mCherry open reading frame (ORF) in pAAV-EF1a-mCherry-DIO (Addgene plasmid #20299, RRID: Addgene_20299) with mt-GFP or 4mt-GCaMP6f (‘mt-GcAMP6f’ in the main text). High-Fidelity Phusion Taq polymerase (Thermo Fisher Scientific, F-530XL) was used to amplify mt-GFP from Addgene plasmid #44385 (RRID: Addgene_44385) or 4mt-GCaMP6f (plasmid was generously donated by Diego de Stefani). For mt-GFP, PCR fragment and plasmid were digested with AscI and NheI (New England Biolabs). Because 4mt-GCaMP6f contains an NheI digestion site, to create the mt-GCaMP6f-DIO plasmid, pAAV-EF1a-mCherry-DIO was digested with NheI, blunted using T4 DNA Polymerase (New England Biolabs), digested with AscI and then dephosphorylated using Antarctic Phosphatase (New England Biolabs). The 4mt-GCamP6f PCR product was phosphorylated using T4 Polynucleotide Kinase (New England Biolabs) and digested using AscI. Plasmid and insert were then ligated overnight at 16 °C (T4 DNA ligase, New England Biolabs). After transformation into chemically competent *E*.*coli* and subsequent plasmid purification, insertion and double inverted orientation (DIO) were confirmed using restriction enzyme analysis and DNA sequencing. The plasmids were deposited to Addgene (mt-GFP-DIO #174112; 4mtGCaMP-DIO: #179529). To test Cre-dependent expression, HEK293-T cells (HEK 293T/17 cell line, CRL-11268 obtained from ATCC; RRID:CVCL_1926) were transfected with pAAV-EF1a-mt-GFP-DIO or pAAV-EF1a-mt-GCaMP6f-DIO only or pAAV-EF1a-mt-GFP-DIO and pCAG*-*Cre (Addgene #13775; RRID Addgene_13775) using polyethylenimine (PEI). Briefly, plasmid DNA (500 ng) was diluted in saline and then mixed with PEI, allowed to incubate for 20-25 minutes and then applied dropwise to HEK293-T cells in a 6-well plate. MitoGFP or mt-GCaMP6f expression was assessed the next day using a fluorescence microscope, which was absent in the cells lacking or expressing only pCAG*-*Cre, confirming Cre-dependence.

mt-GFP primers :

FW: 5’ – AATAGCTAGCATCATGTCCGTCCTGACG – 3’

RV: 5’ - TAATGGCGCGCC GGAAGTCTGGACATTTATTTG – 3’

4mt-GCaMP6f primers:

FW: 5’ – ACT TCG CTG TCA TCA TTT G– 3’

RV: 5’ - GCTAGCCCACCATGTCCGTC– 3’

Sequencing primers:

FW: 5’ – GGATTGTAGCTGCTATTAGC – 3’

RV: 5’ – GGCAAATAACTTCGTATAGGA – 3’

### AAV production

HEK293-T cells of low passage (<25) were kept in Dulbecco′s Modified Eagle′s Medium (DMEM, Thermo Fisher Scientific #31966-047) containing 10% fetal calf serum (FCS, Thermo Fisher Scientific A4766801) and 5% penicillin-streptomycin (Pen-Strep, Thermo Fisher Scientific #15140122) at 37 °C and 5% CO_2_. For virus production, cells were plated on 15 cm dishes at a density of 1-1.25 *×* 10^7^ cells per dish (12 dishes per virus). The next day, medium was replaced with Iscove’s Modified Dulbecco’s Medium (IMDM, Sigma Aldrich I-3390) containing 10% FCS, 5% Pen-Strep and 5% glutamine (Thermo Fisher Scientific #25030081) 1-2 hours before transfection. For transfection, AAV rep/cap, AAV helper (mt-GFP: AAV5; mt-GCaMP6f: AAV1) and transfer plasmids were mixed and diluted in saline before mixing with saline-diluted polyethylenimine (PEI, Polysciences #23966-2) and brief vortexing. After 20-25 minutes of incubation, the transfection mix was added in a drop-wise fashion to the culture plates. The next day, all medium was refreshed after which the cells were left for an additional two nights (*i*.*e*. 72 hours after transfection). Medium was discarded and cells were then collected and subsequently lysed with 3 freeze-thaw cycles to release AAVs. Cell lysate was then loaded on an iodixanol gradient (60, 40, 25 and 15% iodixanol, ELITechGroup #1114542) in Beckman Quick-Seal Polyallomer tubes (Beckman-Coulter #342414) which were sealed and centrifuged in a Beckman-Coulter Optima XE-90 Ultracentrifuge at 16 °C and 69,000 rpm for 1 hour and 10 minutes. The virus-containing fraction was extracted from the tubes and AAVs were then concentrated in Dulbecco’s Phosphate Buffered Saline (D-PBS) + 5% sucrose using Amicon Ultra-15 (100K) filter units (Merck Millipore UFC910024) at 3220 *g*. At least 4 rounds of centrifugation were used to ensure complete replacement of iodixanol with D-PBS + 5% sucrose. Typical yields were ∼150 μl of virus, the titer of which was determined using quantitative PCR (titers: mt-GFP-DIO, 4.43 *×* 10^13^ gc/ml; mt-GCaMP6f-DIO, 3.44 *×* 10^13^ gc/ml)^83^. Viral aliquots were stored at −80 °C until further use.

qPCR primers, recognize AAV2 Inverted Terminal Repeats (ITRs):

5’ – GGAACCCCTAGTGATGGAGTT – 3’

5’ – CGGCCTCAGTGAGCGA – 3’

### Viral injection

Viral injections were typically performed during the 3^rd^ week of control powder food or cuprizone treatment. Mice were anaesthetized using isoflurane (3% induction, 1.2–1.5% maintenance) after which they subcutaneously received 5mg/kg Metacam. Body temperature was monitored and maintained at 37 °C using a heating pad and eyes were prevented from drying out using eye ointment to prevent. The head was then shaved and placed in a stereotaxic frame (Kopf) and an incision was made in the skin along the midline. Lidocaine (10%) was administered on the periost before removing it. Small (<1 mm) bilateral craniotomies were made at -0.5 mm caudally from Bregma and 2.5 mm laterally from the midline without damaging the dura mater. With a sharp glass pipet attached to a Nanoject III (Drummond), 40-50 nL of virus was injected at 1 nL/second and at a depth of 450 μm. Approximately 3 minutes after finishing the injection, the needle was retracted slowly. Bone wax was applied to the craniotomies and the skin was sutured before the animal was allowed to recover.

### Acute slice preparation

After 5-6 weeks of cuprizone or control treatment (*i*.*e*. 2-3 weeks after virus injection), mice were deeply anaesthetized using pentobarbital (50 mg/kg, intraperitoneal injection) and transcranially perfused with ice-cold carbogenated (95% O_2_, 5% CO_2_) cutting artificial cerebrospinal fluid (cACSF; 125 mM NaCl, 3 mM KCl, 6 mM MgCl_2_, 1 mM CaCl_2_, 25 mM glucose, 1.25 mM NaH_2_PO_4_, 1 mM kynurenic acid and 25 mM NaHCO_3_). After quickly dissecting out the brain 400 μm thick parasagittal slices were cut in ice-cold carbogenated cACSF using a Vibratome (1200S, Leica Microsystems). Slices were transferred to a holding chamber containing carbogenated cACSF where they were kept at 35 °C for 35 min to recover and then allowed to return to room temperature for at least 30 min before starting experiments.

### Electrophysiology and two-photon imaging

To perform electrophysiological experiments, slices were transferred to a recording chamber with continuous in- and outflow of carbogenated recording ACSF (rACSF, 125 mM NaCl, 3 mM KCl, 1 mM MgCl_2_, 2 mM CaCl_2_, 10 mM glucose, 5 mM L-Lactate, 1.25 mM NaH_2_PO_4_, and 25 mM NaHCO_3_) at a rate of 1-2 ml per minute at 32 °C. Glass pipettes with an open tip resistance of 6-7 MΩ were filled with an intracellular solution containing (in mM, 130 K-Gluconate, 10 KCl, 10 HEPES, 4 Mg-ATP, 0.3 Na_2_-GTP, 10 Na_2_-phosphocreatine; pH ∼7.25, osmolality ∼280 mOsmol/kg) supplemented with 5 mg/ml biocytin (Sigma-Aldrich, B4261) and 50 μM Atto-594 (Sigma-Aldrich, A08637). Whole-cell recordings were made using a patch-clamp amplifier (Multiclamp 700B, Axon Instruments, Molecular Devices, RRID: SCR_018455) operated by AxoGraph X software (version 1.5.4; RRID: SCR_014284). Action potential trains were evoked using 700 ms step pulses, from −250 pA to +700 nA with increments of 50 pA. Single action potentials were evoked using 3 ms incremental (2.5-5 pA) step pulses with a starting amplitude below spike threshold. Using an AD/DA converter (ITC-18, HEKA Elektronik GmbH), voltage was digitally sampled at 100 Hz. The access resistance during current-clamp experiments (range: 15–30 MΩ) was fully compensated using bridge balance and capacitance neutralization of the amplifier. Somatic single-cell recordings were made from mt-GFP^+^ cells, which were visualized using a two-photon (2P) laser-scanning microscope (Femto3D-RC, Femtonics Inc., Budapest, Hungary). Imaging was controlled using MES software (Femtonics Inc., Budapest, Hungary, version 6.3.7902). A Ti:Sapphire pulsed laser (Chameleon Ultra II, Coherent Inc., Santa Clara, CA, USA) was tuned to 770 nm for two-photon excitation to visualize mt-GFP/mt-GCaMP6f and tdTomato. Fluorescent signals were detected using two photomultipliers (PMTs, Hamamatsu Photonics Co., Hamamatsu, Japan), one for mt-GFP and for Atto-594 (20 μM) to assess its diffusion, as an indication of biocytin diffusion. For motility imaging, *z*-stacks were acquired every 10 s for 8-12 mins. The Image Stabilizer plugin for FIJI was used to account for drift. For Ca^2+^ imaging, mt-GCaMP6f^+^ cells were targeted for single-cell patch clamping and filled with Atto-594 (50 μM) and biocytin (5 mg/ml). Next, *z*-stacks were made with the laser tuned to 800 nm to identify the axon using the Atto-594 signal. Axons were clearly distinguishable from dendrites, as they are much thinner, branch much more often and tend to project back towards the soma, as described previously^20^. Mitochondrial Ca^2+^ responses were then visualized with the laser tuned to 940 nm and recorded at ∼20 Hz imaging frequency. Ca^2+^ responses were analysed using a custom-written Matlab script. Briefly, regions of interest containing mitochondria were selected from which background was subtracted, bleach correction and smoothing were applied, and ΔF/F was calculated. Care was taken to keep the number of APs, which were evoked using a 700 ms block pulse, the same between groups (MBP^−^, 105.6 ± 0.89 action potentials; MBP^+^, 103.6 ± 0.51 APs; cuprizone, 104.7 ± 1.06 APs, Kruskal-Wallis test *P* = 0.1591). MBP^−^ segments do not include AISs.

### Immunohistochemistry

For staining of biocytin-filled cells, upon completion of experiments acute slices were immediately placed into 4% PFA in PBS for 20-25 minutes, followed by 3 washes of 10 minutes with PBS. Next, tissue was blocked for 2 hours in PBS containing 1-2% Triton-X and 10% normal goat serum. After blocking, sections were moved to blocking buffer containing primary antibodies (see **Supplementary Table 1**) and were incubated overnight at room temperature. Following three 10 min washes in PBS, sections were transferred to PBS containing secondary antibodies and were incubated overnight two hours room temperature or overnight at 4 °C. The tissue was finally washed again in PBS three times for 10 minutes before mounting using FluorSave mounting medium (Merck-Millipore #345789). For immunostainings against MBP, the staining protocol was adjusted: blocking was done one hour at 37 °C and one hour room temperature and using 1% Triton, tissue was washed only once in PBS per washing step, and secondary antibodies were incubated one hour at 37 °C and one hour room temperature. For immunostainings on sections without biocytin-filled cells, 400 μm PFA-fixed sections were first cryoprotected by placing them into 30% sucrose-PBS solution until fully saturated. A sliding freezing microtome (Zeiss Hyrax S30; temperature controlled by a Slee Medical GmbH MTR fast cooling unit) was used to cut 40 μm sections, which were either placed in PBS for immediate use or stored at −20 °C in cryoprotectant solution (30% ethylene glycol, 20% glycerol, 0.05 M phosphate buffer) until further use. Immunostaining protocol was the same as for 400 μm sections, but the blocking buffer contained 0.5% Triton-X and incubation duration for secondary antibodies was two hours at room temperature. All steps are performed with gentle shaking, with the exception of the 37 °C incubation steps. For analysis of normal and demyelinated subcortical white matter, Rbp4-Cre mice injected with AAV1-EF1a-mCherry-DIO and AAV5-EF1a-mtGFP-DIO (mixed 1:1) were first perfused with 1x PBS followed by 4% PFA-PBS. Brains were then dissected out and allowed to fix O/N in 4% PFA-PBS after the brains were cryoprotected using 30% sucrose-PBS and processed into 40 μm coronal sections as described above.

### Confocal microscopy

Imaging was performed using a Leica SP5 or SP8 X confocal laser scanning microscope controlled by Leica Application Suite AF. Biocytin-filled cells were imaged with a 63× oil-immersion lens using the tile-scan function with automated sequential acquisition of multiple channels enabled. Step sizes in the Z-axis were 0.3-0.75 μm, and images were collected at a 2048 *×* 2048 pixel resolution at 100-150 Hz. Mitochondria at AISs were imaged similarly, but only cells were imaged whose AIS was directed parallel to the imaging plane, and *z*-stack step sizes were 0.2 μm. Cellular reconstructions and quantification of mitochondrial density and morphology in single cells or at the AIS were performed manually using Neurolucida Software (MBF Bioscience, version 2020.1.3, 64 bit, RRID: SCR_001775). In PV^+^ interneurons, axons could easily be distinguished from dendrites based on their small diameter and extensive branching. Mitochondrial contours were drawn at the plane where the mt-GFP signal was brightest and was omitted when neurite branches deviated strongly from a parallel orientation with respect to the imaging plane (their location was still marked for density calculations). Mitochondria were counted as belonging to the reconstructed cell if the mt-GFP signal was located clearly inside the cytosol and followed the path of the reconstructed neurite. After completion of tracing, reconstructions and contours were analysed using Neurolucida Explorer (MBF Bioscience, 2019.2.1, RRID: SCR_017348), which calculated contour area, aspect ratio (length:width), density, axonal length and inter-mitochondrial distance. For analysis of mitochondrial densities at the myelin sheath, the MBP signal was turned off during tracing of PV^+^ axons and marking their mitochondria, and *vice versa*. FIJI (RRID: SCR_002285) was used to extract partial images for use in figures. After preprocessing in FIJI, Imaris software (Oxford Instruments, version 9.7.2, RRID: SCR_007370) was used to generate three-dimensional surface renderings. The spot functionality of Imaris was used to count mitochondria in the field of view for motility analysis. Motile mitochondria were identified by eye, mitochondria were considered stable if they displaced less than 2 μm.

### 3D Electron microscope data analysis

3D EM data was obtained from the Microns dataset^38^ (www.microns-explorer.org). Basket interneurons were identified based on their morphology, *i*.*e*. their relatively round soma (as opposed to a pyramidal shape), the lack of spines on dendrites and a thin and highly branched axon that was partly myelinated. From each cell, one to two volumes were selected that either contained an axonal branch point with one myelinated and one unmyelinated branch, or a single branch with intermittent myelination. The EM volume of interest and cytosolic segmentation were then downloaded, after which the mitochondria were segmented and the segment length traced manually using Volume Annotation and Segmentation Tool (VAST) software (version 1.4.1)^84^. Because we found that branch points in PV^+^ interneurons are often nodes of Ranvier (a specialized axonal compartment), branch points and mitochondria that resided in them were not included in the analysis. Boutons were excluded for the same reason. If a mitochondrion spanned a border between myelinated and unmyelinated axon or beyond a branch point, they were not included. To analyse the mitochondrial content in L2/3 PNs, we used the same approached as described above. These cells were readily identified based on their pyramidal shape, spiny dendrites and distance from the pia. We selected L2/3 PNs that displayed patchy myelin. Fully or unmyelinated L2/3 PN axons were not included. Length and volume measurements were performed and exported using VAST Tools Matlab scripts. For examples used in figures, 3D models were exported as OBJ files using VAST software and rendered using 3ds Max (Autodesk, version 25.0.0.997, SCR_014251). See **Supplementary Table 2** for cells used in the analysis.

### Statistical analyses

All statistical comparisons were done using Prism (Graphpad, version 8.4.3, RRID: SCR_002798). Normality of datasets was determined using D’Agostino & Pearson or Shapiro-Wilk tests. We applied non-parametric tests if data deviated significantly from a normal distribution. For comparisons between groups we used a two-tailed unpaired t-test (normal data) or a two-tailed Mann-Whitney test (non-normal data). For t-tests degrees of freedom (df) are reported in the figure legends. One-way ANOVA (normal data) or Kruskal-Wallis test (non-normal data) was applied for comparisons of three groups; two-way ANOVA was used to test interactions between groups and treatments; Bonferroni’s post hoc test (normal data) or Dunn’s post hoc test (non-normal data) was used for multiple comparisons; nested t-tests or nested one-way ANOVAs were used when large numbers of datasets were involved (*i*.*e*. mitochondrial contours) to avoid overpowering of non-nested statistical tests (cells were nested inside their respective treatment groups). Figure legends contain *P*-values and *n*-numbers and whether the latter signify mice, cells or mitochondria. Means are presented with SEM.

## Data availability

All data and code are available upon request to the authors.

## Acknowledgments

The authors are thankful to Fred de Winter and Joost Verhaagen (NIN–KNAW) for the support for developing and producing the AAV constructs. We thank Christian Lohmann for critical reading of the manuscript and feedback during this research. We thank Diego de Stefani for generously sharing the 4mt-GCaMP6f plasmid with us. We thank Christiaan Levelt for generously sharing the AAV1-CAG-Flex-mRuby2-GSG-P2A-GCaMP6f virus with us. We thank Arnoldo Zaldivar Castro for optimizing the immunohistochemistry protocol.

## Funding

The Dutch Research Council NWO Vici 865.17.003 (MK)

ZonMW Off Road 04510012010066 (KK)

Erasmus scholarship G ATHINE 01 (MEB)

## Author contributions

Conceptualization: MK, KK

Methodology: MK, KK, MEB, BV

Investigation: KK, MEB, BV, NP

Formal analysis, MK, KK, MEB, BV

Visualization: MK, KK

Funding acquisition: MK

Project administration: MK

Supervision: MK, KK

Writing – original draft: KK

Writing – review & editing: MK, KK

## Supplementary Figures

**Supplementary Figure 1.**
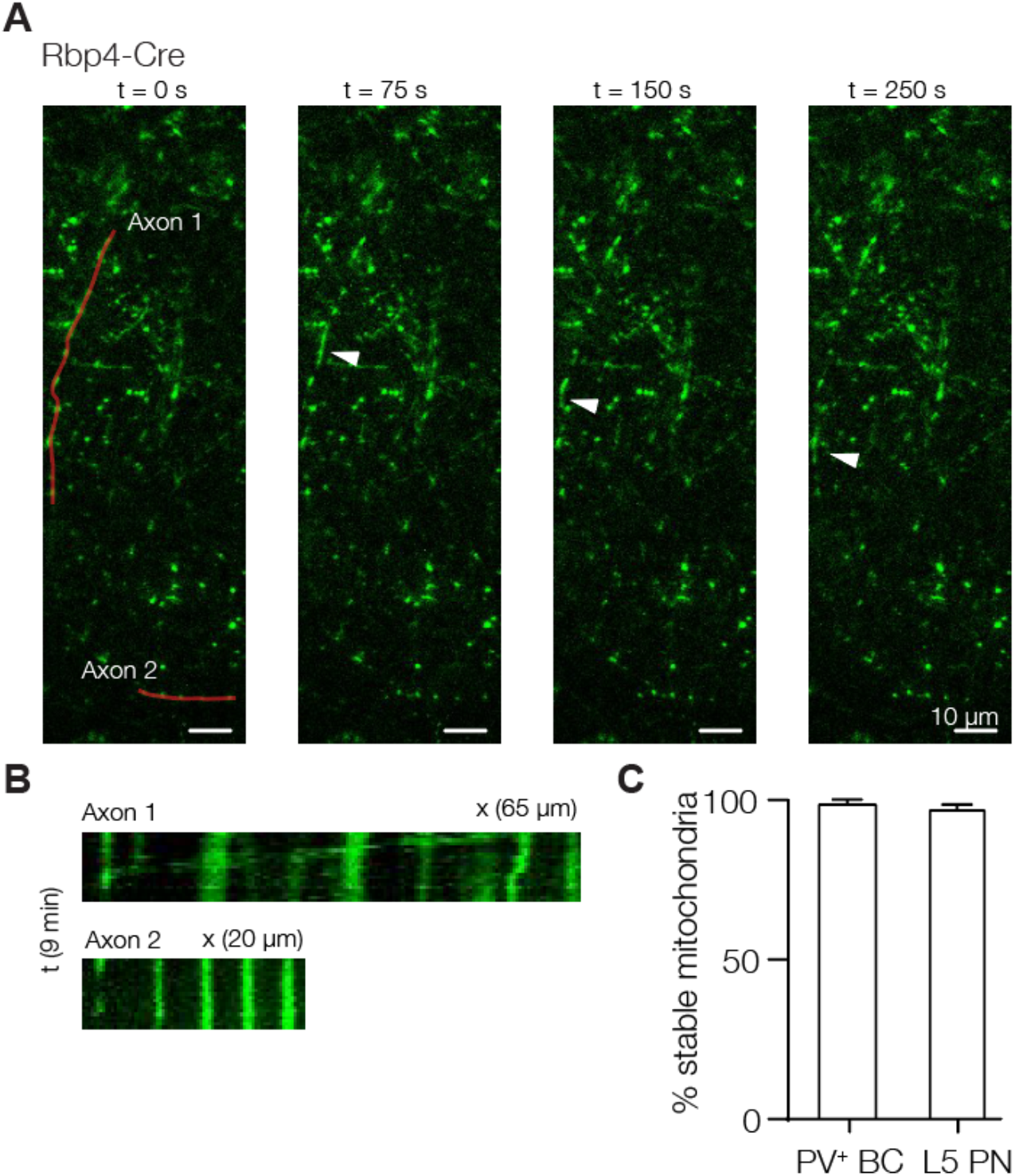
Mitochondrial motility in acute brain slices. **(A)** Example frames showing motile mitochondria (mt-GFP, green) in one L5 PN axon, but all other mitochondria are stationary. Red lines indicate paths used for kymographs in b. **(B)** Example kymographs showing mitochondrial motility (1) and stability (2). **(C)** Quantification of stationary mitochondria in PV^+^ interneurons (PV^+^ basket cells (BC); PV-Cre; Ai14 mice, *n =* 4) or L5 PNs (Rbp4-Cre mice, *n =* 3). PV-Cre; Ai14, *n =* 11/1236; Rbp4-Cre, *n =* 23/1145 motile mitochondria/total mitochondria.

**Supplementary Figure 2.**
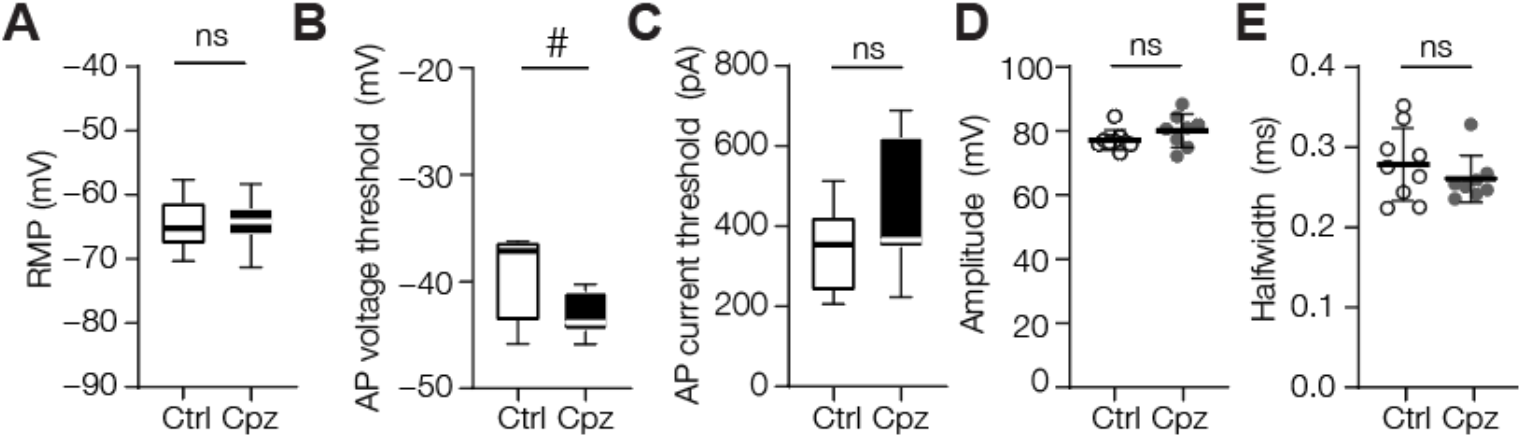
Action potential parameters and mitochondrial distribution at the AIS upon demyelination. **(A)** No change in resting membrane potential (RMP; unpaired t-test, *P =* 0.7249; control, *n =* 9 cells from 5 mice; cuprizone, *n =* 8 cells from 5 mice). **(B)** Voltage threshold shows a trend towards reduction upon cuprizone-mediated demyelination (unpaired t-test, *P =* 0.0590; control, *n =* 9 cells from 5 mice; cuprizone, *n =* 8 cells from 5 mice); No difference between control and cuprizone-treated groups in **(C)** current threshold (unpaired t-test, *P =* 0.2066), **(D)** AP amplitude (unpaired t-test, 0.1697) or **(E)** halfwidth (unpaired t-test, *P =* 0.3489; control, *n =* 9 cells from 5 mice; cuprizone, *n =* 8 cells from 5 mice). Horizontal bars indicate the mean, individual data points represent cells, error bars indicate SEM; Box plots represent the 25^th^ to 75^th^ percentiles, whiskers indicate the maximal and minimal values, solid line represents the median.

**Supplementary Figure 3.**
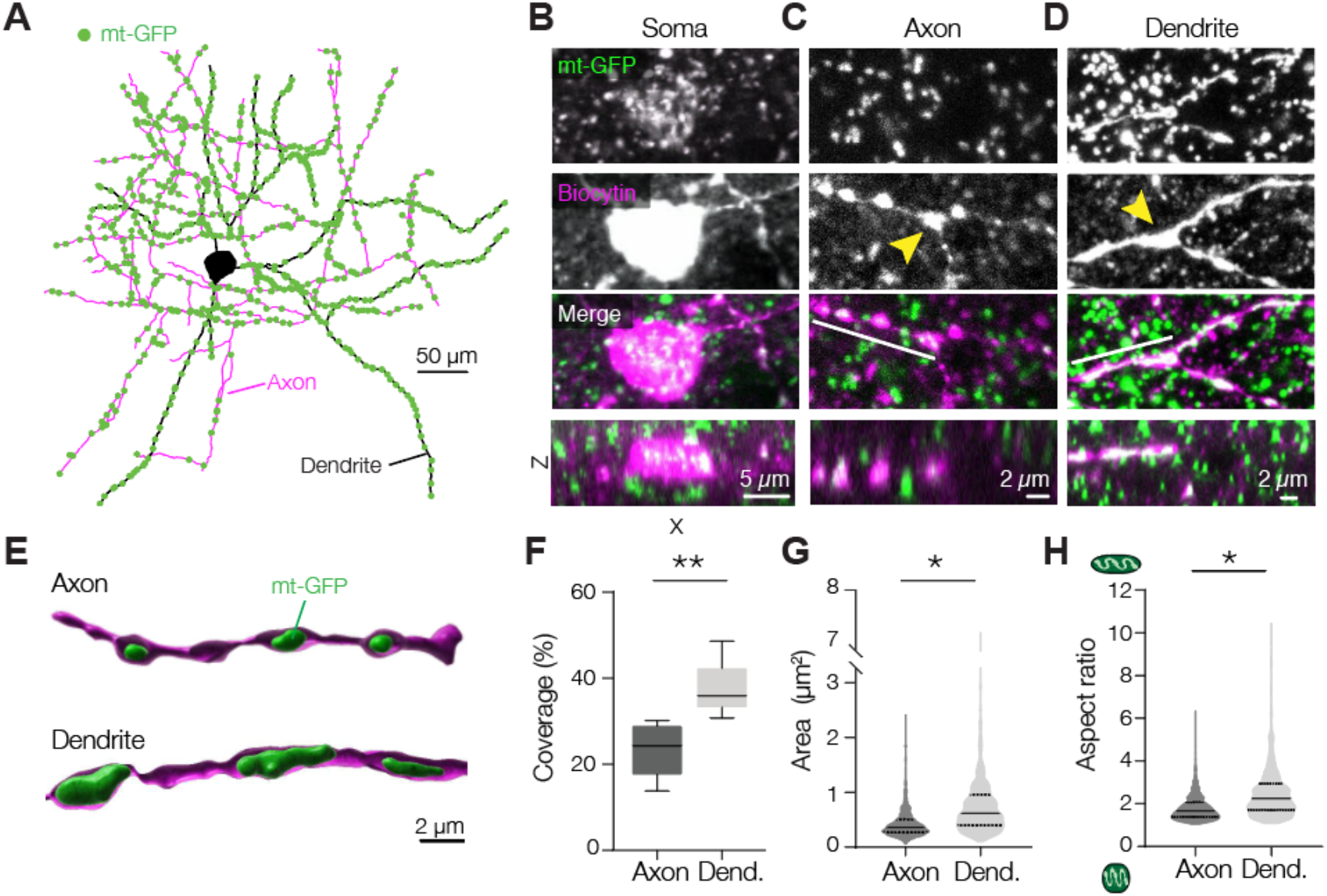
Axonal and dendritic PV^+^ interneuron mitochondria are morphologically distinct. **(A)** Example of a reconstructed PV^+^ interneuron and the locations of its mitochondria (green). Magenta, axon; black, dendrite. Mitochondria in soma were not quantified. **(B)** Examples of somatic, **(C)** axonal and **(D)** dendritic mitochondria in PV^+^ interneurons. Yellow arrowheads indicate branch points, white lines indicate axis used for *XZ* views. **(E)** 3D surface rendering of example axonal or dendritic mitochondria. **(F)** Percentage of PV^+^ neurite length covered by mitochondrial length (unpaired t-test, \*\**P =* 0.0093; df = 8; Axon, *n =* 188 mitochondria; dendrite, *n =* 482 mitochondria; 5 cells from 3 mice). **(G)** Area of mitochondria is larger in PV^+^ dendrites (nested t-test, \**P =* 0.0144; df = 8; Axon, *n =* 859 mitochondria; dendrite, *n =* 1138 mitochondria; 5 cells from 3 mice). **(H)** Aspect ratio of mitochondria is higher in PV^+^ dendrites, reflecting a more elongated morphology (nested t-test, \**P =* 0.0318; df = 8; Axon, *n =* 859 mitochondria; dendrite, *n =* 1138 mitochondria; 5 cells from 3 mice). Box plots (f-h) represent the 25^th^ to 75^th^ percentiles, whiskers represent the maximal and minimal values, solid line represents the median. Solid lines in truncated violin plots represent the median, dotted lines represent 25^th^ and 75^th^ quartiles.

**Supplementary Figure 4.**
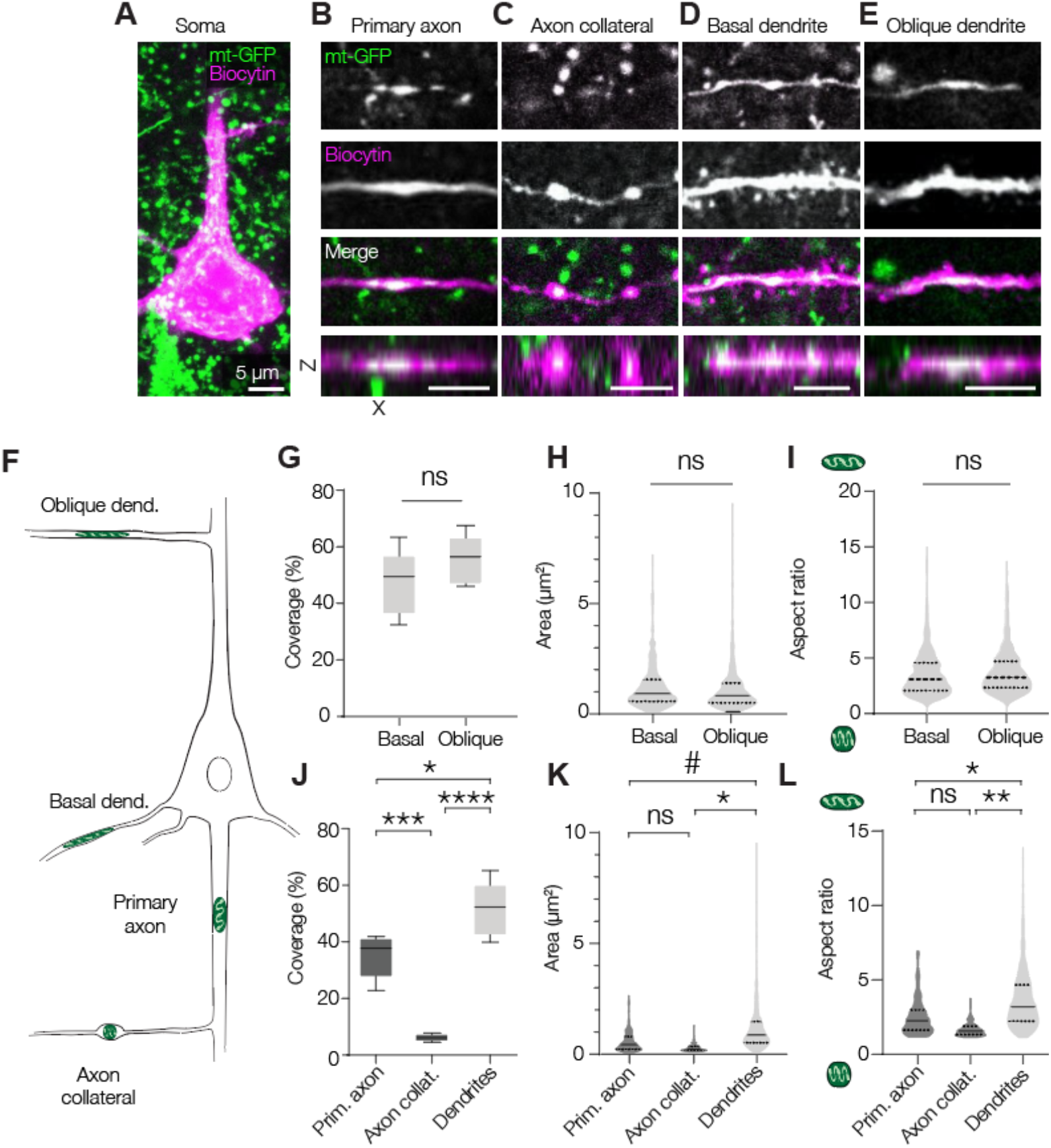
Axonal mitochondria are distinct from those in dendrites in L5 PNs. Example images of mitoGFP-labeled mitochondria in the **(A)** soma, **(B)** primary axon, **(C)** axon collateral, **(D)** oblique dendrite branch or **(E)** basal dendrite of a L5 PN. Somatic mitochondria were not quantified. **(F)** Schematic of subcellular compartments of the L5 PN that were investigated. **(G)** Dendritic coverage is not different between basal and oblique dendrites (unpaired t-test, *P =* 0.2538; basal dendrites, *n =* 321; oblique dendrites, *n =* 331 mitochondria from 5 cells of 3 mice each for each). **(H)** Mitochondria from basal and oblique dendrites have similar surface area (nested t-test, *P =* 0.9554) and **(I)** aspect ratio (nested t-test, *P =* 0.7162; basal dendrites, *n =* 321; oblique dendrites, *n =* 331 mitochondria from 5 cells of 3 mice each for each). **(J)** Neurite length covered by mitochondrial length (one-way ANOVA, *P* < 0.0001; Bonferroni’s post hoc test, Primary axon vs axon collaterals, \*\*\**P =* 0.0001; primary axon vs dendrites, \*\**P =* 0.0141; axon collateral vs dendrites, \*\*\*\**P* < 0.0001; primary axon *n =* 188, axon collateral *n =* 125, dendrites *n =* 652 mitochondria; for each, *n =* 5 cells from 3 mice). **(K)** Mitochondrial area in L5 PNs is largest in dendrites and smallest in axon collaterals (nested one-way ANOVA, \**P =* 0.0192; Bonferroni’s post hoc test, primary axon vs axon collaterals *P* > 0.9999; primary axon vs dendrites, #*P =* 0.0917; axon collaterals vs dendrites, \**P =* 0.0241; primary axon *n =* 188, axon collateral *n =* 125, dendrites *n =* 652 mitochondria; for each, *n =* 5 cells from 3 mice). **(L)** Mitochondria are significantly more round in L5 PN axons compared to dendrites (nested one-way ANOVA, *P =* 0.0031; Bonferroni’s post hoc test, primary axon vs axon collaterals, *P =* 0.5094; primary axon vs dendrites, \*\**P =* 0.0418; axon collaterals vs dendrites, \*\*\**P =* 0.0031; primary axon *n =* 188, axon collateral *n =* 125, dendrites *n =* 652 mitochondria; for each, *n =* 5 cells from 3 mice). Box plots represent the 25^th^ to 75^th^ percentiles, whiskers represent the maximal and minimal values, solid line represents the median. In truncated violin plots, solid lines represent the median, dotted lines represent 25^th^ and 75^th^ quartiles.

**Supplementary Figure 5.**
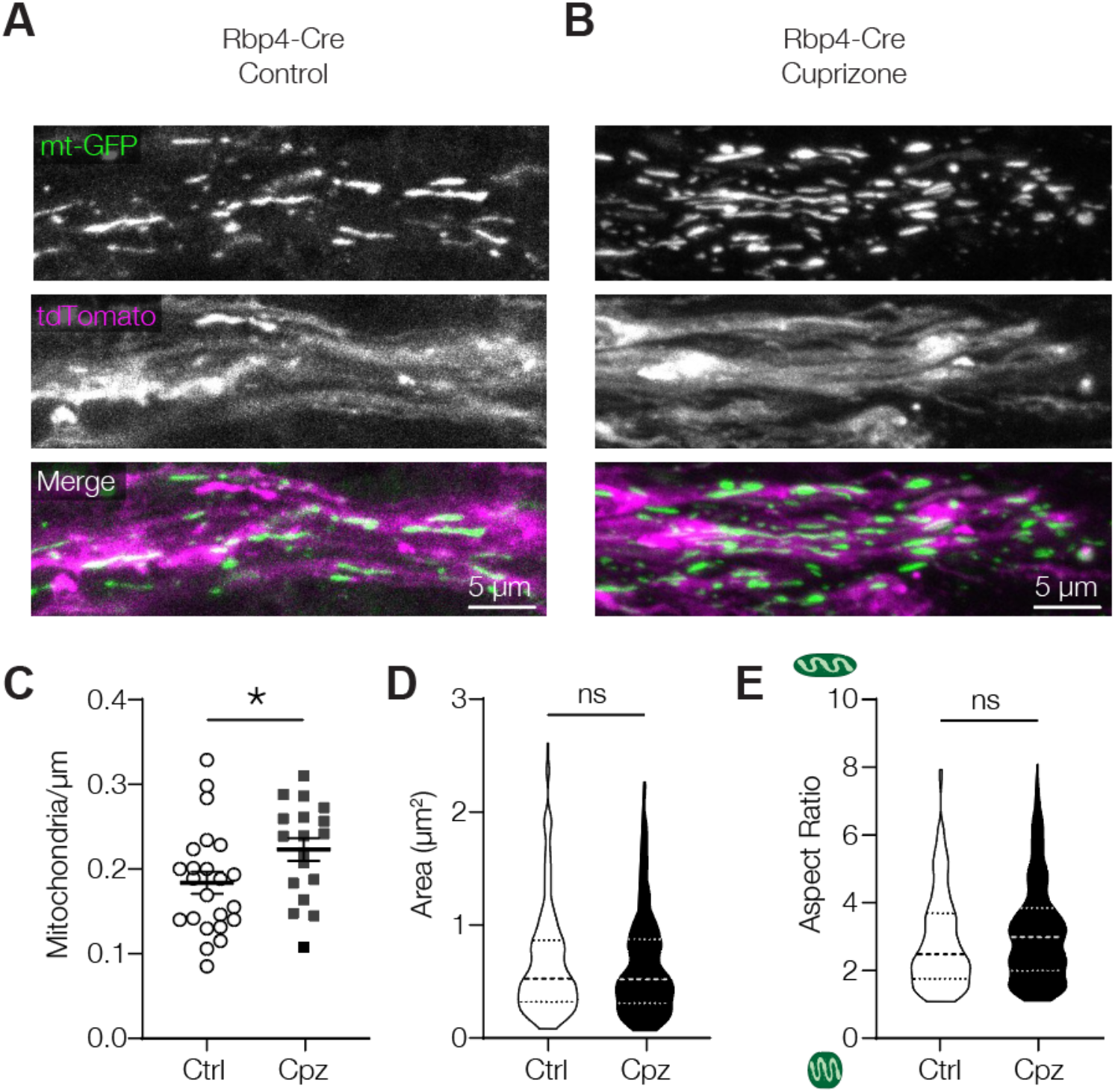
Increased mitochondrial density in the demyelinated white matter excitatory axons. **(A-B)** Example confocal images of mitochondria in the white matter of control (A) or cuprizone-treated (B) L5 pyramidal neuron axons from Rbp4-Cre mice. **(C)** Increased density after cuprizone-mediated demyelination (unpaired t-test, \**P* = 0.0441; control, *n* = 23 axons from 3 mice; cuprizone, *n* = 19 axons from 3 mice). **(D)** Mitochondrial size is unchanged after demyelination (nested t-test, *P* = 0.6886; control, *n* = 172 mitochondria from 23 axons of 3 mice; cuprizone, *n* = 162 mitochondria from 19 axons of 3 mice). **(E)** Comparable aspect ratio of mitochondria in control or cuprizone-treated white matter axons (nested t-test, *P* = 0.1668; control, *n* = 172 mitochondria from 23 axons of 3 mice; cuprizone, *n* = 162 mitochondria from 19 axons of 3 mice). Horizontal lines in c the mean, error bars represent SEM, individual data points indicate axons. In truncated violin plots, solid lines represent the median, dotted lines represent 25^th^ and 75^th^ quartiles.

**Supplementary Figure 6.**
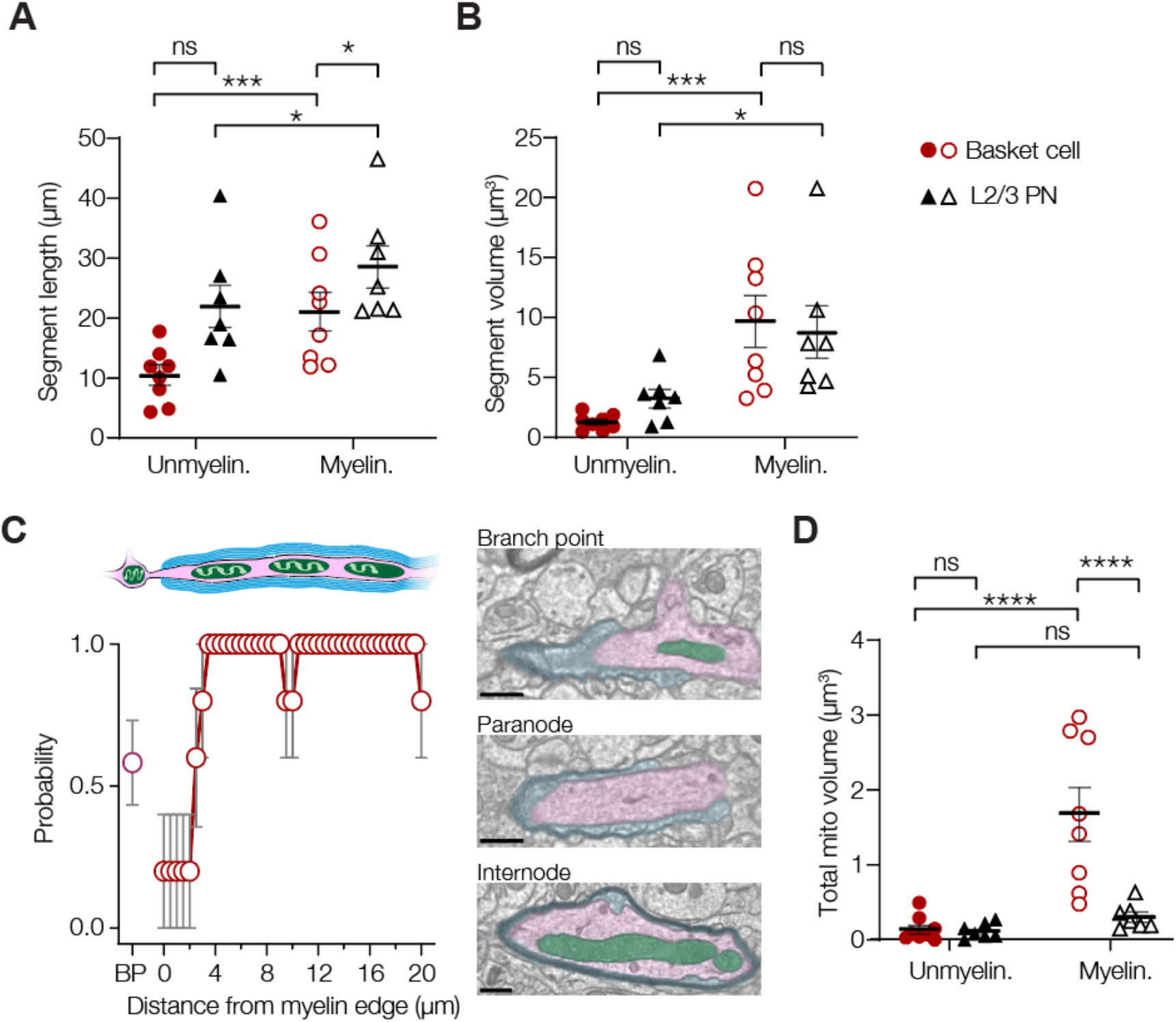
Myelination-dependent internode properties and mitochondria distribution. **(A)** Segment length of (un)myelinated axons in basket cells or L2/3 PNs (two-way ANOVA, myelination x cell type effect, *P =* 0.2139, F_(1,13)_ = 1.708; myelination effect, *P* < 0.0001, F_(1,13)_ = 32.78; cell type effect, *P =* 0.0339, F_(1,13)_ = 5.619; Bonferroni’s post hoc test, basket myelinated vs. unmyelinated, \*\*\**P* = 0.0004; PN myelinated vs. unmyelinated, \**P* = 0.0195; Myelinated basket vs PN, *P =* 0.1804; Unmyelinated basket vs PN \**P* = 0.0253). **(B)** Segment volume of (un)myelinated axons in basket cells or L2/3 PNs (two-way ANOVA, myelination x cell type effect, *P =* 0.2765, F_(1,13)_ = 1.209; myelination effect, *P* = 0.0001, F_(1,13)_ = 28.30; cell type effect, *P =* 0.7825, F_(1,13)_ = 0.07945; Bonferroni’s post hoc test, basket myelinated vs. unmyelinated, \*\*\**P* = 0.0008; PN myelinated vs. unmyelinated, \**P* = 0.0265; Myelinated basket vs PN, *P >* 0.9999; Unmyelinated basket vs PN *P* = 0.7632). **(C)** Probability of mitochondria occupancy at branch points (BP; presumed nodes) and along the myelinated internode. *Left:* Mitochondria are often found at branch points (probability 0.58; 12 branch points from 8 cells), show limited occupancy in the first ∼2 μm of the internode and are distributed evenly and with high probability along the internode (5 internodes from 4 cells). *Right:* example EM images of a branchpoint containing a mitochondrion, a paranode devoid of mitochondria and an internode containing a large mitochondrion. Axons, myelin and mitochondria are pseudocoloured (magenta, blue and green, respectively). Scale bar, 500 nm. **(D)** Total volume of mitochondria in (un)myelinated axons in basket cells or L2/3 PNs (two-way ANOVA, myelination x cell type effect, *P =* 0.0027, F_(1,13)_ = 13.58; myelination effect, *P* = 0.0004, F_(1,13)_ = 22.16; cell type effect, *P =* 0.0052, F_(1,13)_ = 11.24; Bonferroni’s post hoc test, basket myelinated vs. unmyelinated, \*\*\*\**P <* 0.0001; PN myelinated vs. unmyelinated, *P =* 0.9925; Myelinated basket vs PN, \*\*\*\**P <* 0.0001; Unmyelinated basket vs PN *P* > 0.9999).

**Supplementary Figure 7.**
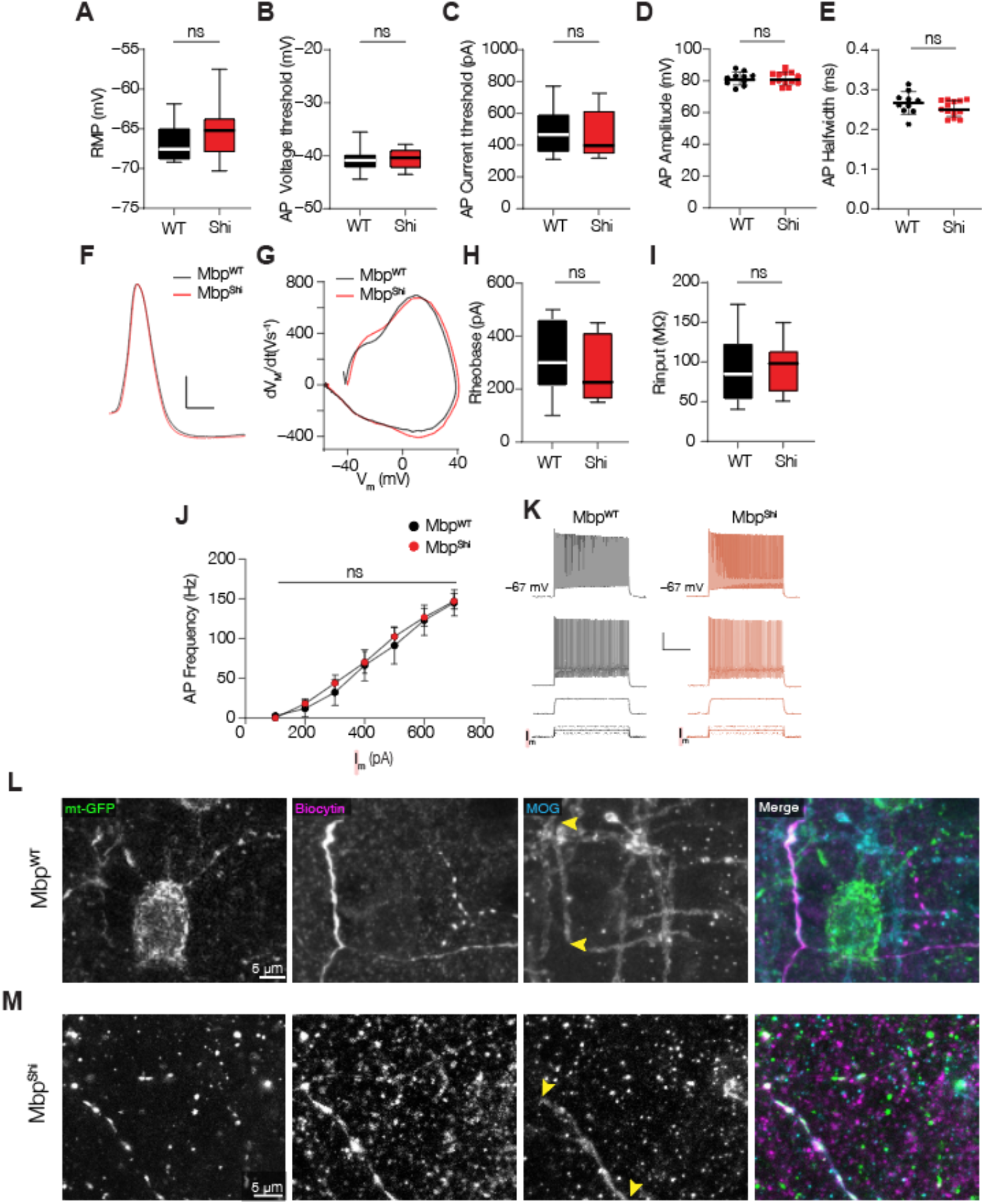
Electrophysiological properties of PV^+^ interneurons in *Mbp*^*WT*^ or *Mbp*^*Shi*^ mice. **(A)** Resting membrane potential (unpaired t-test, *P* = 0.2597; df = 18; *Mbp*^*WT*^, *n* = 8 cells from 3 mice; *Mbp*^*Shi*^, *n* = 12 cells from 3 mice); **(B)** Action potential membrane threshold (unpaired t-test, *P* = 0.8364; df = 21; *Mbp*^*WT*^, *n* = 10 cells from 3 mice; *Mbp*^*Shi*^, *n* = 13 cells from 3 mice); **(C)** Action potential current threshold (unpaired t-test, *P* = 0.6566; df = 21; *Mbp*^*WT*^, *n* = 10 cells from 3 mice; *Mbp*^*Shi*^, *n* = 13 cells from 3 mice); **(D)** Action potential amplitude (unpaired t-test, *P* = 0.9358; df = 21; *Mbp*^*WT*^, *n* = 10 cells from 3 mice; *Mbp*^*Shi*^, *n* = 13 cells from 3 mice); **(E)** Action potential halfwidth (unpaired t-test, *P* = 0.1744; df = 21; *Mbp*^*WT*^, *n* = 10 cells from 3 mice; *Mbp*^*Shi*^, *n* = 13 cells from 3 mice); **(F)** Example single action potential from either genotype, scalebar indicates 10 mV and 0.25 ms; **(G)** Phase-plane plots of the example traces in h; **(H)** Rheobase (unpaired t-test, *P* = 0.4395; df = 18; *Mbp*^*WT*^, *n* = 8 cells from 3 mice; *Mbp*^*Shi*^, *n* = 12 cells from 3 mice); **(I)** Input resistance (unpaired t-test, *P* = 0.8375; df = 18; *Mbp*^*WT*^, *n* = 8 cells from 3 mice; *Mbp*^*Shi*^, *n* = 12 cells from 3 mice); **(J)** Comparable action potential firing frequency in *Mbp*^*WT*^ or *Mbp*^*Shi*^ mice (two-way ANOVA, *P =* 0.9967, F_(13, 221)_ = 0.2488; genotype effect *P =* 0.7752, F_(1, 17)_ = 0.08422; *Mbp*^*WT*^, *n* = 8 cells from 3 mice; *Mbp*^*Shi*^, *n* = 12 cells from 3 mice); **(K)** Example traces in response to 200, 400 or 600 pA somatic current injection, scale bar indicates 25 mV and 250 ms.). **(L-M)** Confocal images corresponding to the 3D renders in Figure 6. Yellow arrowheads indicate internodes. Individual data points in a-k represent cells, error bars indicate SEM; Box plots represent the 25th to 75th percentiles, whiskers indicate the maximal and minimal values, solid line represents the median.

**Supplementary Figure 8.**
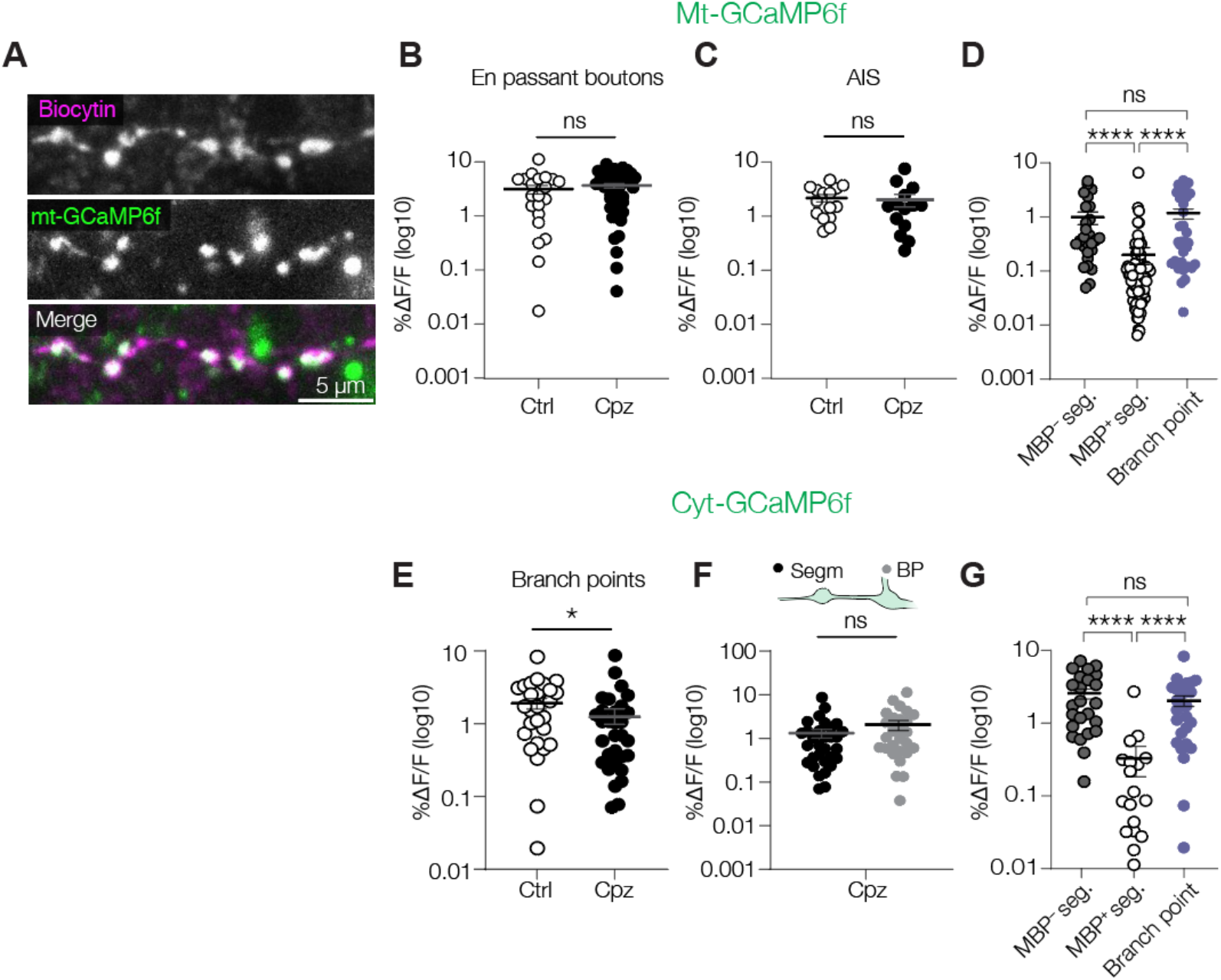
mt-Ca^2+^ and cyt-Ca^2+^ responses in PV^+^ axons. **(A)** Example confocal image of PV^+^ *en passant* boutons. **(B)** Demyelination does not alter mt-Ca^2+^ buffering in *en passant* boutons (Mann-Whitney test, *P* = 0.2847; control, *n =* 21 mitochondria of 4 cells from 2 mice; cuprizone, *n* = 47 mitochondria of 8 cells from 4 mice). **(C)** mt-Ca^2+^ responses at the AIS are not changed after demyelination (Mann-Whitney test, *P* = 0.1694; control, *n* = 15 mitochondria of 5 AISs from 3 mice; cuprizone, *n* = 20 mitochondria of 6 AISs from 4 mice). **(D)** In control PV^+^ axons, amplitude of mt-Ca^2+^ responses at MBP^−^ segments are not different from those at branch points (Kruskal-Wallis test, *P* < 0.0001; Dunn’s post hoc test, MBP^+^ vs MBP^−^ *P* = 0.0020, MBP^−^ vs. branch points *P* > 0.9999; MBP^+^ vs branch points *P* < 0.0001; MBP^+^ segments, *n =* 101 mitochondria of 12 cells from 7 mice, same data as in Figure 7D; MBP^−^ segments, *n* = 27 mitochondria of 9 cells from 5 mice, same data as in Figure 7D; branch points, *n* = 38 mitochondria of 11 cells from 7 mice, same data as in Figure 8D); **(E)** Smaller cyt-Ca^2+^ transients in branch points after demyelination (Mann-Whitney test, *P* = 0.0307; control, *n* = 27 branch points of 7 cells from 3 mice; cuprizone, *n* = 31 branch points of 6 cells from 3 mice); **(F)** Cytosolic Ca^2+^ responses are comparable in segments and branch points upon demyelination (Mann-Whitney test, *P* = 0.2476; *n* = 28 segments of 6 cells from 3 mice, *n* = 31 branch points of 6 cells from 3 mice, same data as in E); **(G)** In control PV^+^ axons, amplitude of cytosolic Ca^2+^ responses at MBP^−^ segments are not different from those at branch points (Kruskal-Wallis test, *P* < 0.0001; Dunn’s post hoc test, MBP^+^ vs MBP^−^ *P* < 0.0001, MBP^−^ vs. branch points *P* > 0.9999; MBP^+^ vs branch points *P* < 0.0001; MBP^+^, *n* = 18 segments of 6 cells from 3 mice; MBP^−^, *n* = 24 segments of 8 cells from 3 mice, same data as in Figure 7F; branch points, control, *n* = 27 branch points of 7 cells from 3 mice, same data as in E).

**Supplementary Table 1.**

| **Antibody** | **Host** | **Dilution** | **Manufacturer** | **Catalog number** | **RRID** |
| --- | --- | --- | --- | --- | --- |
| Anti-Green fluorescent protein | Chicken | 1:500 | Abcam | ab13970 | AB_300798 |
| Anti-Red fluorescent protein | Mouse | 1:500 | Abcam | ab65856<br>(discontinued) | AB_1141717 |
| Anti-βIV Spectrin | Rabbit | 1:1000 | Engelhardt and Kole lab | N/A | N/A |
| Anti-Myelin Basic Protein | Mouse | 1:250 | Covance | SMI-99P | AB_10120129 |
| Anti-Myelin Oligodendrocyte Glycoprotein | Mouse | 1:250 | Millipore | MAB5680 | RRID:AB_1587278 |
| Anti-Mouse-Alexa-405 | Goat | 1:500 | ThermoFisher | A31553 | AB_221604 |
| Anti-Chicken-Alexa-488 | Goat | 1:500 | ThermoFisher | A11039 | AB_142924 |
| Anti-Rabbit-Alexa-633 | Goat | 1:500 | ThermoFisher | A21070 | AB_2535731 |
| Anti-Mouse-Alexa-647 | Goat | 1:500 | ThermoFisher | A21235 | AB_2535804 |
| Streptavidin-Alexa-594 | N/A | 1:500 | ThermoFisher | S11227 | N/A |
| Streptavidin-Alexa-633 | N/A | 1:500 | ThermoFisher | S32364 | AB_2313500 |

Overview of antibodies and dilutions used in this study.

**Supplementary Table 2.**

| Cell name | Neuroglancer ID | Cell type | Location of interest |
| --- | --- | --- | --- |
| PV1_1 | 864691135307240262 | Basket cell | 174508, 124118, 21157 |
| PV1_2 | 864691135307240262 | Basket cell | 180056, 128599, 21514 |
| PV2_1 | 864691135367308281 | Basket cell | 178363, 187957, 20579 |
| PV3_1 | 864691135386593409 | Basket cell | 181574, 159259, 20999 |
| **PV3_2** | **864691135386593409** | **Basket cell** | **191862, 145103, 20142** |
| PV4_1 | 864691135544579880 | Basket cell | 171765, 173397, 20706 |
| PV5_1 | 864691135428608048 | Basket cell | 181752, 143942, 22968 |
| PV6_1 | 864691135590150923 | Basket cell | 170564, 132985, 19749 |
| PV7_1 | 864691135697462549 | Basket cell | 173603, 134391, 22067 |
| PV8_1 | 864691136952074207 | Basket cell | 186366, 118464, 20183 |
| PN_1 | 864691135855890478 | L2/3 PN | 144874, 149452, 23959 |
| PN_2 | 864691135975539779 | L2/3 PN | 337852, 147425, 19387 |
| PN_3 | 864691135564739159 | L2/3 PN | 334986, 157943, 20515 |
| PN_4 | 864691136031786427 | L2/3 PN | 123284, 152471, 23565 |
| PN_5 | 864691136618414989 | L2/3 PN | 222968, 137995, 23424 |
| PN_6 | 864691135012580342 | L2/3 PN | 238983, 137249, 23366 |
| **PN_7** | **864691135099882016** | **L2/3 PN** | **241318, 147339, 23370** |

Overview of the cells used in the 3D EM analysis. Bold marked cells were used as examples in **Figure 5**.

## Supplementary Videos

### Supplementary Video 1

Heatmap of mitochondrial Ca^2+^ responses in a PV^+^ axons upon 100 APs. A significant response can be seen in the branch point but not under the myelin sheath. Video corresponds to Figure 7b.

### Supplementary Video 1

Heatmap of cytosolic Ca^2+^ responses in a PV^+^ axons upon 100 APs. In the AIS and the first branch point, significant responses are seen but not under the myelinated internode. Video corresponds to Figure 7e.

## References

1. Cohen, C. C. H. et al. Saltatory Conduction along Myelinated Axons Involves a Periaxonal Nanocircuit. Cell 180, 311-322.e15 (2020).

2. Nave, K.-A. & Werner, H. B. Myelination of the Nervous System: Mechanisms and Functions. Annu Rev Cell Dev Bi 30, 503–533 (2014).

3. Lee, Y. et al. Oligodendroglia metabolically support axons and contribute to neurodegeneration. Nature 487, 443–448 (2012).

4. Fünfschilling, U. et al. Glycolytic oligodendrocytes maintain myelin and long-term axonal integrity. Nature 485, 517–521 (2012).

5. Chamberlain, K. A. et al. Oligodendrocytes enhance axonal energy metabolism by deacetylation of mitochondrial proteins through transcellular delivery of SIRT2. Neuron (2021) doi:10.1016/j.neuron.2021.08.011.

6. Philips, T. et al. MCT1 Deletion in Oligodendrocyte Lineage Cells Causes Late-Onset Hypomyelination and Axonal Degeneration. Cell Reports 34, 108610 (2021).

7. Zambonin, J. L. et al. Increased mitochondrial content in remyelinated axons: implications for multiple sclerosis. Brain 134, 1901–1913 (2011).

8. Witte, M. E. et al. Enhanced number and activity of mitochondria in multiple sclerosis lesions. J Pathology 219, 193–204 (2009).

9. Mahad, D. J. et al. Mitochondrial changes within axons in multiple sclerosis. Brain 132, 1161–1174 (2009).

10. Kiryu-Seo, S., Ohno, N., Kidd, G. J., Komuro, H. & Trapp, B. D. Demyelination increases axonal stationary mitochondrial size and the speed of axonal mitochondrial transport. J Neurosci 30, 6658--66 (2010).

11. Nikic, I. et al. A reversible form of axon damage in experimental autoimmune encephalomyelitis and multiple sclerosis. Nat Med 17, 495--499 (2011).

12. Licht-Mayer, S. et al. Enhanced axonal response of mitochondria to demyelination offers neuroprotection: implications for multiple sclerosis. Acta Neuropathol 140, 143–167 (2020).

13. Ohno, N. et al. Mitochondrial immobilization mediated by syntaphilin facilitates survival of demyelinated axons. Proc National Acad Sci 111, 9953–9958 (2014).

14. Licht-Mayer, S. et al. Axonal response of mitochondria to demyelination and complex IV activity within demyelinated axons in experimental models of multiple sclerosis. Neuropath Appl Neuro e12851 (2022) doi:10.1111/nan.12851.

15. Andrews, H. et al. Increased axonal mitochondrial activity as an adaptation to myelin deficiency in the Shiverer mouse. J Neurosci Res 83, 1533–1539 (2006).

16. Balaratnasingam, C., Morgan, W. H., Johnstone, V., Cringle, S. J. & Yu, D.-Y. Heterogeneous Distribution of Axonal Cytoskeleton Proteins in the Human Optic Nerve. Invest Ophth Vis Sci 50, 2824–2838 (2009).

17. Perge, J. A., Koch, K., Miller, R., Sterling, P. & Balasubramanian, V. How the Optic Nerve Allocates Space, Energy Capacity, and Information. J Neurosci 29, 7917–7928 (2009).

18. Kageyama, G. H. & Wong-Riley, M. T. The histochemical localization of cytochrome oxidase in the retina and lateral geniculate nucleus of the ferret, cat, and monkey, with particular reference to retinal mosaics and ON/OFF-center visual channels. J Neurosci 4, 2445–59 (1984).

19. Kasthuri, N. et al. Saturated Reconstruction of a Volume of Neocortex. Cell 162, 648–661 (2015).

20. Stedehouder, J. et al. Fast-spiking Parvalbumin Interneurons are Frequently Myelinated in the Cerebral Cortex of Mice and Humans. Cereb Cortex 27, 5001–5013 (2017).

21. Micheva, K. D. et al. A large fraction of neocortical myelin ensheathes axons of local inhibitory neurons. Elife 5, e15784 (2016).

22. Dubey, M. et al. Myelination synchronizes cortical oscillations by consolidating parvalbumin-mediated phasic inhibition. Elife 11, (2022).

23. Kepecs, A. & Fishell, G. Interneuron cell types are fit to function. Nature 505, 318--26 (2014).

24. Hu, H., Gan, J. & Jonas, P. Fast-spiking, parvalbumin<sup>+</sup> GABAergic interneurons: From cellular design to microcircuit function. Science 345, 1255263 (2014).

25. Stedehouder, J. et al. Local axonal morphology guides the topography of interneuron myelination in mouse and human neocortex. Elife 8, e48615 (2019).

26. Edgar, J. M. et al. Early ultrastructural defects of axons and axon–glia junctions in mice lacking expression of Cnp1. Glia 57, 1815–1824 (2009).

27. Evangelou, N. et al. Size-selective neuronal changes in the anterior optic pathways suggest a differential susceptibility to injury in multiple sclerosis. Brain 124, 1813–1820 (2001).

28. Ramaglia, V. et al. Complement-associated loss of CA2 inhibitory synapses in the demyelinated hippocampus impairs memory. Acta Neuropathol 142, 643–667 (2021).

29. Dutta, R. et al. Mitochondrial dysfunction as a cause of axonal degeneration in multiple sclerosis patients. Ann Neurol 59, 478--489 (2006).

30. Zoupi, L. et al. Selective vulnerability of inhibitory networks in multiple sclerosis. Acta Neuropathol 141, 415–429 (2021).

31. Hu, H., Roth, F. C., Vandael, D. & Jonas, P. Complementary Tuning of Na+ and K+ Channel Gating Underlies Fast and Energy-Efficient Action Potentials in GABAergic Interneuron Axons. Neuron 98, 156-165.e6 (2018).

32. Li, T. et al. Action Potential Initiation in Neocortical Inhibitory Interneurons. Plos Biol 12, e1001944 (2014).

33. Kann, O., Papageorgiou, I. E. & Draguhn, A. Highly Energized Inhibitory Interneurons are a Central Element for Information Processing in Cortical Networks. J Cereb Blood Flow Metabolism 34, 1270–1282 (2014).

34. Kageyama, G. H. & Wong-Riley, M. T. T. Histochemical localization of cytochrome oxidase in the hippocampus: Correlation with specific neuronal types and afferent pathways. Neuroscience 7, 2337–2361 (1982).

35. Kontou, G. et al. Miro1-dependent mitochondrial dynamics in parvalbumin Interneurons. Elife 10, e65215 (2021).

36. Stedehouder, J. & Kushner, S. A. Myelination of parvalbumin interneurons: a parsimonious locus of pathophysiological convergence in schizophrenia. Mol Psychiatr 22, 4–12 (2017).

37. Micheva, K. D. et al. Distinctive Structural and Molecular Features of Myelinated Inhibitory Axons in Human Neocortex. Eneuro 5, ENEURO.0297-18.2018 (2018).

38. Consortium, Mic. et al. Functional connectomics spanning multiple areas of mouse visual cortex. Biorxiv 2021.07.28.454025 (2021) doi:10.1101/2021.07.28.454025.

39. Fecher, C. et al. Cell-type-specific profiling of brain mitochondria reveals functional and molecular diversity. Nat Neurosci 22, 1731--1742 (2019).

40. Meyer, H. S. et al. Inhibitory interneurons in a cortical column form hot zones of inhibition in layers 2 and 5A. Proc National Acad Sci 108, 16807--12 (2011).

41. Gerfen, C. R., Paletzki, R. & Heintz, N. GENSAT BAC Cre-Recombinase Driver Lines to Study the Functional Organization of Cerebral Cortical and Basal Ganglia Circuits. Neuron 80, 1368–1383 (2013).

42. Hamada, M. S., Goethals, S., Vries, S. I. D., Brette, R. & Kole, M. H. P. Covariation of axon initial segment location and dendritic tree normalizes the somatic action potential. Proc National Acad Sci 113, 14841--14846 (2016).

43. Hamada, M. S. & Kole, M. H. P. Myelin Loss and Axonal Ion Channel Adaptations Associated with Gray Matter Neuronal Hyperexcitability. J Neurosci 35, 7272–7286 (2015).

44. Smit-Rigter, L. et al. Mitochondrial Dynamics in Visual Cortex Are Limited In Vivo and Not Affected by Axonal Structural Plasticity. Curr Biol 26, 2609–2616 (2016).

45. Kipp, M., Clarner, T., Dang, J., Copray, S. & Beyer, C. The cuprizone animal model: new insights into an old story. Acta Neuropathol 118, 723–736 (2009).

46. Kole, M. H. P., Letzkus, J. J. & Stuart, G. J. Axon Initial Segment Kv1 Channels Control Axonal Action Potential Waveform and Synaptic Efficacy. Neuron 55, 633–647 (2007).

47. Kole, M. H. & Brette, R. The electrical significance of axon location diversity. Curr Opin Neurobiol 51, 52–59 (2018).

48. Zilberter, Y., Zilberter, T. & Bregestovski, P. Neuronal activity in vitro and the in vivo reality: the role of energy homeostasis. Trends Pharmacol Sci 31, 394–401 (2010).

49. Cserep, C., Posfai, B., Schwarcz, A. D. & Denes, A. Mitochondrial ultrastructure is coupled to synaptic performance at axonal release sites. Eneuro 5, (2018).

50. Lewis, T. L., Kwon, S.-K., Lee, A., Shaw, R. & Polleux, F. MFF-dependent mitochondrial fission regulates presynaptic release and axon branching by limiting axonal mitochondria size. Nat Commun 9, 5008 (2018).

51. Tomassy, G. S. et al. Distinct Profiles of Myelin Distribution Along Single Axons of Pyramidal Neurons in the Neocortex. Science 344, 319–324 (2014).

52. Edgar, J. M., McCulloch, M. C., Thomson, C. E. & Griffiths, I. R. Distribution of mitochondria along small-diameter myelinated central nervous system axons. J Neurosci Res 86, 2250–2257 (2008).

53. Saab, A. S. & Nave, K.-A. Myelin dynamics: protecting and shaping neuronal functions. Curr Opin Neurobiol 47, 104–112 (2017).

54. Roach, A., Takahashi, N., Pravtcheva, D., Ruddle, F. & Hood, L. Chromosomal mapping of mouse myelin basic protein gene and structure and transcription of the partially deleted gene in shiverer mutant mice. Cell 42, 149–155 (1985).

55. Snaidero, N. et al. Myelin Membrane Wrapping of CNS Axons by PI(3,4,5)P3-Dependent Polarized Growth at the Inner Tongue. Cell 156, 277–290 (2014).

56. Hanemaaijer, N. A. et al. Ca2+ entry through NaV channels generates submillisecond axonal Ca2+ signaling. Elife 9, e54566 (2020).

57. Zhang, Z. & David, G. Stimulation-induced Ca2+ influx at nodes of Ranvier in mouse peripheral motor axons. J Physiology 594, 39–57 (2016).

58. Ashrafi, G., Juan-Sanz, J. de, Farrell, R. J. & Ryan, T. A. Molecular Tuning of the Axonal Mitochondrial Ca2+ Uniporter Ensures Metabolic Flexibility of Neurotransmission. Neuron 105, 678-687.e5 (2020).

59. Stoler, O. et al. Frequency- and spike-timing-dependent mitochondrial Ca2+ signaling regulates the metabolic rate and synaptic efficacy in cortical neurons. Elife 11, e74606 (2022).

60. Crawford, D. K., Mangiardi, M., Xia, X., López-Valdés, H. E. & Tiwari-Woodruff, S. K. Functional recovery of callosal axons following demyelination: a critical window. Neuroscience 164, 1407–1421 (2009).

61. Dupree, J. L. et al. Oligodendrocytes assist in the maintenance of sodium channel clusters independent of the myelin sheath. Neuron Glia Biol 1, 179–192 (2004).

62. Micheva, K. D., Kiraly, M., Perez, M. M. & Madison, D. V. Conduction Velocity Along the Local Axons of Parvalbumin Interneurons Correlates With the Degree of Axonal Myelination. Cereb Cortex 31, bhab018. (2021).

63. Wortman, J. C. et al. Axonal Transport: How High Microtubule Density Can Compensate for Boundary Effects in Small-Caliber Axons. Biophys J 106, 813–823 (2014).

64. Griffiths, I. et al. Axonal Swellings and Degeneration in Mice Lacking the Major Proteolipid of Myelin. Science 280, 1610–1613 (1998).

65. Mazuir, E., Fricker, D. & Sol-Foulon, N. Neuron–Oligodendrocyte Communication in Myelination of Cortical GABAergic Cells. Life 11, 216 (2021).

66. Pekkurnaz, G., Trinidad, J. C., Wang, X., Kong, D. & Schwarz, T. L. Glucose Regulates Mitochondrial Motility via Milton Modification by O-GlcNAc Transferase. Cell 158, 54–68 (2014).

67. Mensch, S. et al. Synaptic vesicle release regulates myelin sheath number of individual oligodendrocytes in vivo. Nat Neurosci 18, 628–630 (2015).

68. Almeida, R. G. et al. Myelination induces axonal hotspots of synaptic vesicle fusion that promote sheath growth. Curr Biol (2021) doi:10.1016/j.cub.2021.06.036.

69. Rangaraju, V., Calloway, N. & Ryan, T. A. Activity-Driven Local ATP Synthesis Is Required for Synaptic Function. Cell 156, 825–835 (2014).

70. Young, E. A. et al. Imaging correlates of decreased axonal Na+/K+ ATPase in chronic multiple sclerosis lesions. Ann Neurol 63, 428–435 (2008).

71. Gennerich, A. & Vale, R. D. Walking the walk: how kinesin and dynein coordinate their steps. Curr Opin Cell Biol 21, 59–67 (2009).

72. Mironov, S. L. ADP Regulates Movements of Mitochondria in Neurons. Biophys J 92, 2944–2952 (2007).

73. Turner, N. L. et al. Reconstruction of neocortex: Organelles, compartments, cells, circuits, and activity. Cell 185, 1082-1100.e24 (2022).

74. Benamer, N., Vidal, M., Balia, M. & Angulo, M. C. Myelination of parvalbumin interneurons shapes the function of cortical sensory inhibitory circuits. Nat Commun 11, 5151 (2020).

75. Grundemann, J. & Clark, B. A. Calcium-Activated Potassium Channels at Nodes of Ranvier Secure Axonal Spike Propagation. Cell Reports 12, 1715--1722 (2015).

76. Popovic, M. A., Foust, A. J., McCormick, D. A. & Zecevic, D. The spatio-temporal characteristics of action potential initiation in layer 5 pyramidal neurons: a voltage imaging study. J Physiology 589, 4167–4187 (2011).

77. Foust, A. J., Yu, Y., Popovic, M., Zecevic, D. & McCormick, D. A. Somatic Membrane Potential and Kv1 Channels Control Spike Repolarization in Cortical Axon Collaterals and Presynaptic Boutons. J Neurosci 31, 15490–15498 (2011).

78. Rizzuto, R., Stefani, D. D., Raffaello, A. & Mammucari, C. Mitochondria as sensors and regulators of calcium signalling. Nat Rev Mol Cell Bio 13, 566–578 (2012).

79. Trevisiol, A. et al. Monitoring ATP dynamics in electrically active white matter tracts. Elife 6, e24241 (2017).

80. Stedehouder, J., Brizee, D., Shpak, G. & Kushner, S. A. Activity-Dependent Myelination of Parvalbumin Interneurons Mediated by Axonal Morphological Plasticity. J Neurosci 38, 3631–3642 (2018).

81. Yang, S. M., Michel, K., Jokhi, V., Nedivi, E. & Arlotta, P. Neuron class–specific responses govern adaptive myelin remodeling in the neocortex. Science 370, (2020).

82. Persson, A.-K., Hoeijmakers, J. G. J., Estacion, M., Black, J. A. & Waxman, S. G. Sodium Channels, Mitochondria, and Axonal Degeneration in Peripheral Neuropathy. Trends Mol Med 22, 377–390 (2016).

83. Aurnhammer, C. et al. Universal Real-Time PCR for the Detection and Quantification of Adeno-Associated Virus Serotype 2-Derived Inverted Terminal Repeat Sequences. Hum Gene Ther Part B Methods 23, 18–28 (2012).

84. Berger, D. R., Seung, H. S. & Lichtman, J. W. VAST (Volume Annotation and Segmentation Tool): Efficient Manual and Semi-Automatic Labeling of Large 3D Image Stacks. Front Neural Circuit 12, 88 (2018).

